# BayesRB: a markov chain Monte Carlo-based polygenic genetic risk score algorithm for dichotomous traits

**DOI:** 10.1101/2022.02.27.482193

**Authors:** Ying Shan, Daniel E. Weeks

## Abstract

Identifying high-risk individuals with diseases through reliable prediction models guides screening and preventive treatment. Most complex diseases have a genetic basis influenced by multiple genes and so disease risk can be estimated using polygenic risk score (PRS) algorithms. Many PRS algorithms have been developed so far. Among them, BayesR shows good characteristics of unbiasedness, accuracy, sparseness, and robustness. It detects the associated SNPs, estimates the SNP effects, and makes prediction of disease risks based on all SNPs simultaneously. However, this method assumes that the phenotypes follow a Gaussian distribution, which cannot be met in case-control studies. Here, we made an extension of the BayesR method, called BayesRB, by adding auxiliary variables to the BayesR model. We explored the characteristics, efficacy, and accuracy of BayesRB when estimating SNP effects and predicting disease risks compared with three traditional algorithms under different conditions using both simulated data and real data from the Welcome Trust Case Control Consortium (WTCCC). For SNP effect estimation, BayesRB shows unbiasedness and sparseness for big and small effect SNPs, respectively. For disease risk prediction, BayesRB had the best performance among the methods. This study provides a theoretical basis for complex disease risk prediction and disease prevention

## 1 Introduction

Disease prevention is the most economical and effective health strategy [Khera et al., 2018]. The key to prevention lies in the disease risk prediction and identification of high-risk individuals [Lander, 2011]. Most common diseases have a genetic component [Fuchsberger et al., 2016; Golan et al., 2014], so disease risk prediction can benefit from the full use of genetic information [Eeltink et al., 2021]. Therefore, polygenic risk score (PRS) algorithms have been developed for complex disease prediction. Using PRS algorithms, a large number of genetic markers spanning the whole genome can be integrated, so as to comprehensively predict individual disease risk using a genetic basis [Lewis and Vassos, 2020; Li et al., 2020; Choi et al., 2020a].

Most polygenic risk scores (PRS) are constructed using single nucleotide polymorphism (SNP) markers. The “threshold method” and the “shrinkage method” are two common strategies [Choi et al., 2020b]. The “threshold method” clumps and thins the SNPs to make them independent of each other, selects SNPs with p values less than the threshold, and uses the estimated SNP effect sizes as weights to construct an additive model. The “threshold method” is commonly used, but it has a series of defects, including (1) SNP selection is subjective, due to the use of a subjective selection threshold; (2) The SNP effects are marginal effects, and the failure to account for the effects of other SNPs in the linkage disequilibrium (LD) block result in large error variance, and low risk prediction power [Moser et al., 2015]; (3) Considering multiple testing in genome-wide association studies (GWAS), stringent P-value thresholds are often used - SNPs which are significant under a stringent threshold may only account for a small part of heritability, thus leading to low risk prediction power. (4) When using the selected threshold, the “winner’s curse” will inflate the effect size estimates of the selected variants, thus leading to suboptimal genetic predictions [Huang et al., 2018]. The “shrinkage method” shrinks the estimated effect size of whole-genome SNP by statistical methods and builds the PRS models with the estimated effect size after shrinkage. One of the most commonly used “shrinkage methods”, the penalized generalized linear model, shrinks the weights of SNPs with small effects to 0 by applying L1 regularization or a combination of L1 and L2 regularization across the genome. The penalized generalized linear regression models assume that each SNP effect follows the same distribution with a common variance [Chiaia et al., 2017] This assumption is strong and can limit the biological interpretability of results [Mollandin et al., 2021].

A series of Bayesian models (known as the Bayesian alphabet) have been developed with varying models for the distribution of SNP effects. In BayesA, all SNP effects are assumed to be drawn from a normal distribution whose variance follows a scale inverse *χ*^2^ prior distribution [Meuwissen et al., 2001]. BayesB and BayesC assume that a fixed portion of markers have zero effect and the variance of non-null SNPs respectively follow a per-SNP or common inverse *χ*^2^ distribution [Mollandin et al., 2021]. BayesC*π* and BayesD*π* assume that the proportion of null SNP effects *π* is a random variable [Habier et al., 2011]. BayesR models SNP effects with a four-component normal mixture model, thus it is more flexible than previous Bayesian models, allowing some SNPs to have zero effect while others to have small to moderate effects. BayesR was shown to yield accurate predictions and promise for quantitative trait loci (QTL) mapping [Mollandin et al., 2021].

On the basis of BayesR, many extensions were developed. EmBayesR uses the expectation-maximization (EM) algorithm to replace the Markov chain Monte Carlo (MCMC). EmBayesR provides similar estimates of SNP effects and accuracies of genomic prediction to bayesR, but reduces computing time [Wang et al., 2015]. BayesMV takes into account the situation that one polymorphism affects multiple traits, and can be used for simultaneously elucidating genetic architecture, QTL mapping, and genomic prediction [Kemper et al., 2018]. LDAK-BayesR-Predict and LDAK-BayesR-SS models allow the users to assume an effect size distribution that is a mixture of a point mass at zero and three Gaussian distributions [Zhang et al., 2021]. SBayes extends bayesR to the one that uses GWAS summary statistics [Lloyd-Jones et al., 2019]. Besides the estimation of SNP effects and the point estimates of traits, Bayesian PRS models are able to estimate the variance of an individual’s PRS and obtain well-calibrated credible intervals by using posterior sampling [Ding et al., 2021].

Most BayesR extended models are based on the assumption that the phenotypes are continuous and are not directly applicable to dichotomous phenotypes encountered in case/control studies. While BayesR can be applied by the coding the binary outcome as 0/1, this may cause the following problems: 1) when fitting an ordinary linear model, the residuals do not follow normal distributions, but instead follow a standard logistic distribution, which may cause biased SNP effect estimation and biased risk prediction; 2) the SNP odds ratios cannot be calculated using the estimated SNP effects; 3) it may be hard to explain the predicted outcome: the predicted phenotypes can be bigger than 1, so it cannot be treated as the probability of being a case.

Several previous studies have proposed methods for dealing with dichotomous outcome data [Ma and Zhou, 2021]. Directly fitting of a Bayesian logistic model is a method that can be used. However, one drawback of fitting a Bayesian logistic model directly is that there is no conjugate prior. Most previous approaches applied Metropolis-Hastings, or otherwise used accept-reject steps [Chen and Dey, 1998; Gamerman, 1997]. These make each MCMC step complicated and computationally intensive. When analyzing hundreds of thousands of SNPs, it is necessary that each MCMC step runs quickly. Simple Gibbs sampling can be used when the auxiliary variable models are used. Albert and Chib [1993] proposed an approximate data-augmentation algorithm, using *t*(8) quantiles to approximate the logistic quantiles. The posterior distributions of all the parameters have standard forms. Thus, Gibbs sampling can be applied. But this is an approximate method, instead of an exact one. Polson et al. [2013] proposed another data-augmentation strategy for Bayesian logistic regression, which is a direct analogue of Albert and Chib’s construction. Their approach is based on a newly proposed Polya-Gamma distribution family. But it is not practical in a Bayesian mixture model. Holmes and Held [2006] proposed an exact data-augmentation algorithm. In their approach, one posterior distribution does not have a standard form so they used rejection sampling to solve the issue. However, adaptive-rejection sampling only updates individual coefficients, which leads to a poor mixing when coefficients are correlated. Therefore, they suggested a joint updating of some parameters whose posteriors correlated to each other. This allows fast mixing in the chain.

Here, we developed a Bayesian method called BayesRB for dichotomized phenotypes, which was an extension of BayesR based on Erbe et al. [2012] and Moser et al [2015]. BayesRB selects associated SNPs, estimates the SNP effects, and makes the prediction of the disease risks, while taking into account for the effects of SNPs in the whole genome. We also compared the performance of BayesRB with “threshold methods”, lasso, and BayesR using both simulation datasets and real datasets. BayesRB was shown to be a promising method for dichotomized disease/traits prediction.

## 2 Methods

### 2.1 BayesRB Approach

To estimate each SNP effect taking account for all the SNP effects, we constructed a Bayesian mixture model and assumed the SNP effects come from mixtures of several normal distributions, including one with zero mean and zero variance. We applied the Markov chain Monte Carlo (MCMC) to estimate the unknown parameters. We used a Gibbs scheme and Metropolis-Hasting scheme to sample values from each unknown parameter’s conditional posterior distribution. Then, we made inference of each unknown parameters, including the effect sizes of the SNPs and the categories the SNPs belongs to.

In a sample with *n* independent individuals and *p* independent SNPs, the phenotypes are related to SNPs with a logistic regression model

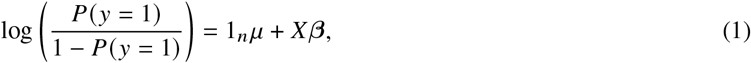

where *y* is an *n*-dimensional vector of dichotomous phenotypes, *P*(*y* = 1) is the probability of being affected, 1_*n*_ is an *n*-dimensional vector of ones, *μ* is the general mean of the *P*(*y* = 1) in the logit scale, *X* is a *n* × *p* matrix of genotypes coded as 0, 1, or 2 indicating 0, 1, or 2 risk alleles. The vector ***β*** is a *p*-dimensional vector of SNP effects.

To extend the BayesR approach to binary traits, we introduced an auxiliary variable *Z_i_*.

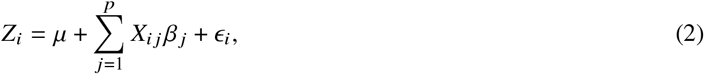

where *ϵ_i_* follows a standard logistic distribution. For the SNP *j*, ***X_.j_*** is standardized with the mean of 0 and the variance of 1, where ***X_.j_*** is the vector of the number of risk alleles of SNP *j* for all the individuals. To keep the conditional conjugacy for updating *β*, we introduced a further set of variables, **λ**, which contains *λ_i_*, *i* = 1, …, *n*. Then, the model becomes

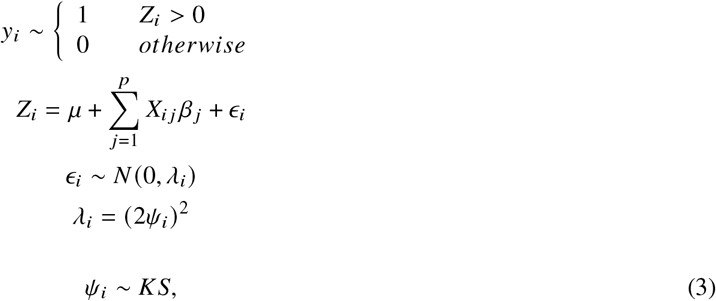

where *ψ_i_, i* = 1,…, *n*, are independent variables following the Kolmogorov-Smirnov (KS) distribution. This model is equivalent to the logistic regression model [Holmes and Held, 2006]. The *SNP_j_*’s effect *β_j_* is assumed to be a mixture of four zero mean normal distributions.

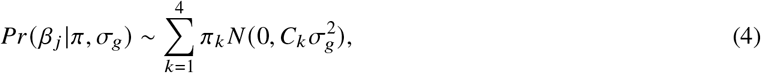

where *k* = (1, 2, 3, 4). (*C*_1_, *C*_2_, *C*_3_, *C*_4_) = (0, 10^−4^, 10^−3^, 10^−2^). 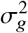 is the genetic variance. There are two strategies to deal with 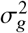, according to Erbe et al. and Moser et al., respectively: 1) treating 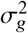 as fixed; 2) treating 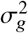 as random. When treating 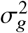 as random, we set 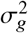 follows a uniform non-informative prior with the initial value follows *N*(0, 200). ***π*** = (*π*_1_, *π*_2_, *π*_3_, *π*_4_) are the mixture proportions, which sum up to 1. The prior for *π* is a symmetric Dirichlet distribution:

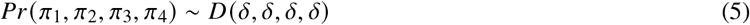

with *δ* = 1.

For the MCMC process, the fully conditional posterior distributions of each unknown parameters are given below. The proof can be found in the Appendix. We use |. to represent conditioned on the data and all other parameters. The dependency diagram treating 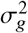 as fixed can be found in Figure 1, and treating 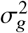 as random can be found in Figure 2.

1. Update {***Z*** and *λ*} jointly from their joint conditional posterior distribution:

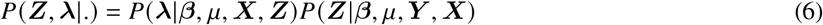 In this case, the ***Z***|***β**, μ, **Y**, **X*** follows independent truncated logistic distribution as shown below:

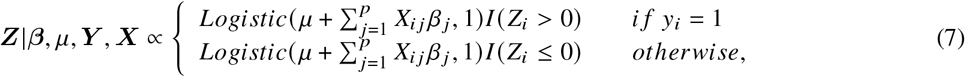

where *Logistic*(*θ*, *κ*) is the density function of the logistic distribution with a mean of *θ* and a scale of *κ*. The conditional distribution *P*(**λ**|***β**, μ, **X**, **Z***) does not have a standard form. Therefore, we used a rejection sampling process to sample the *λ*’s.
2. The conditional posterior distribution of the general mean *μ* is

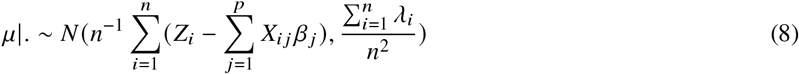

*μ* is sampled from the above distribution for each cycle of the Markov chain.
3. For SNP *j*, *β_j_* and *b_j_* are updated jointly from their joint conditional posterior distribution, where ***b*** denotes the category that each SNP belongs to. SNP *j* belongs to cetegory *k* (k=1, 2, 3, or 4): *b_j_* = *k*.

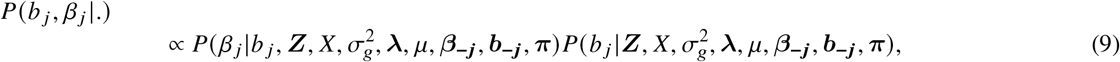

where ***b_−j_*** denotes the vector of categories that all the SNPs expect SNP *j* belong to. ***β_−j_*** denotes the vector of effects of all the SNPs expect SNP *j*. ***C*** is the vector (0, 10^−4^, 10^−3^, 10^−2^). Then,

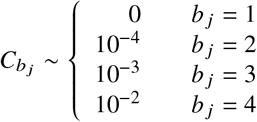 For each SNP, *b_j_* is updated first. Set 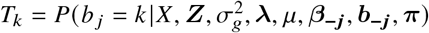.

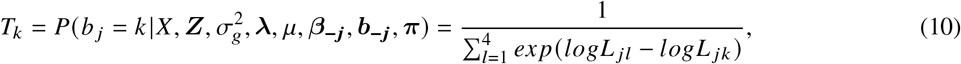

where *k* is the category SNP *j* is assigned to. *k* = (1, 2, 3, 4). If *k* ≠ 1,

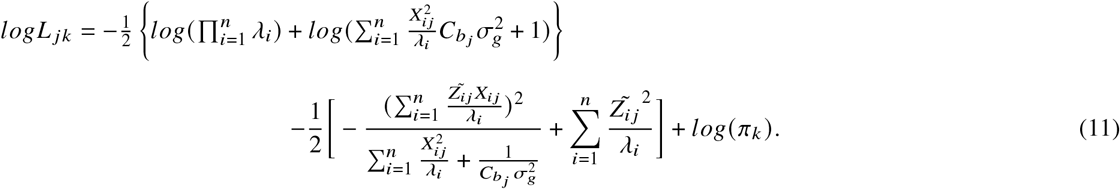 If *k* = 1,

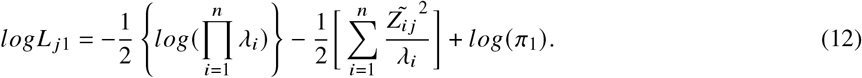 The SNP *j* is assigned to category *k* based on a value *h* sampled from a uniform distribution.

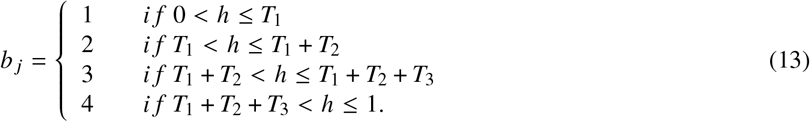 Then, we updated *β_j_*:

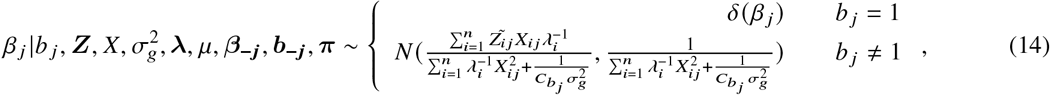

where 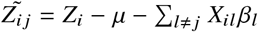, and *δ*(*β_j_*) denotes the dirac delta function with all probability mass at *β_j_* = 0 if *b_j_* = 1. For each cycle of the Markov chain, *b_j_* and *β_j_* for SNP *j* are updated using the above distribution. Then, we repeated step 3 for SNP *j* + 1,…, *p*, and recorded the number of SNPs being in each category as *m_k_*. *k* = (1, 2, 3, 4).
4. If treating 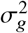 as fixed, then skip this step. If estimating the 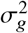, then we set a uniform prior for 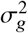 and updated it using the Metropolis-Hasting sampling for each cycle of the Markov chain. We set the truncated normal distribution with variance of *θ* as the proposal function. The detail can be found in the Appendix.
5. The conditional posterior distribution of the mixture proportion *π* follows

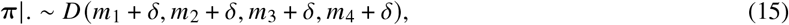

where *δ* = 1. And ***π*** is sampled from the above distribution for each cycle of the Markov chain.
6. Following Moser et al. [2015], in order to increase mixing, we randomly permuted the SNP orders. Then, loop back to step 1.

**Figure 1:**
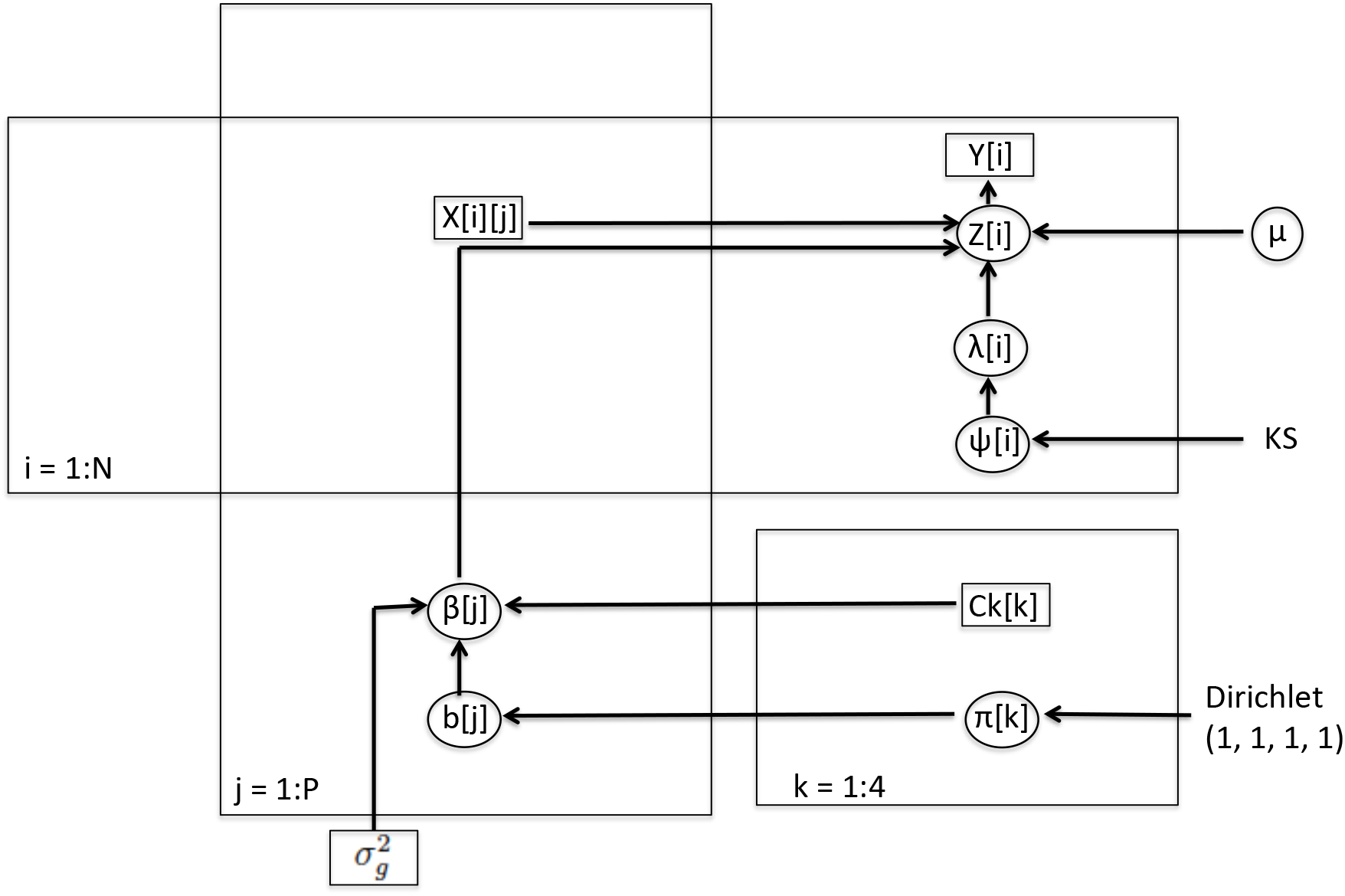
Dependency diagram of the parameters if treating 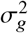 as fixed.

**Figure 2:**
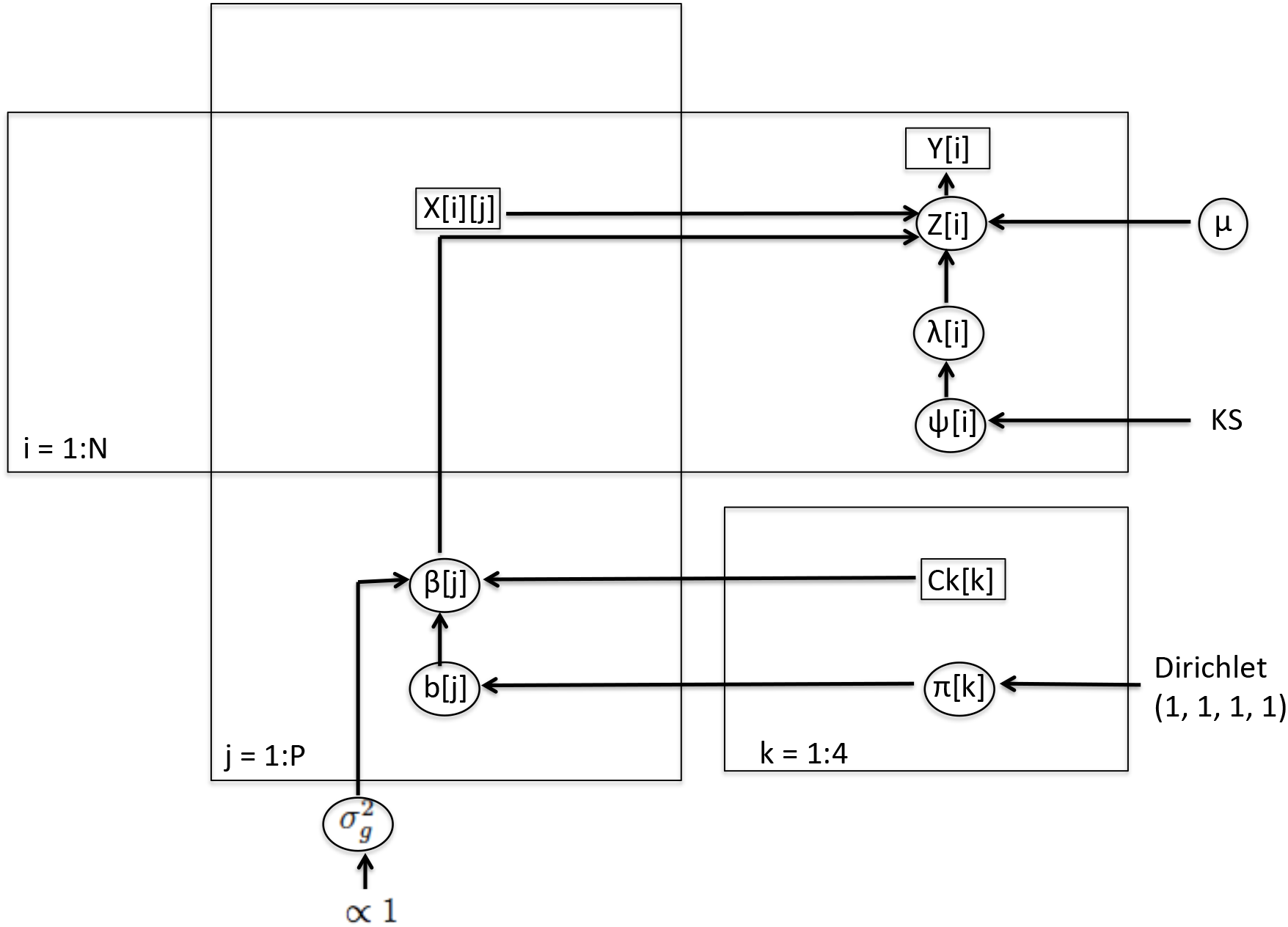
Dependency diagram of the parameters if treating 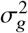 as random.

The pseudo code of the above MCMC steps can be found in the Appendix.

After the MCMC steps, the parameters can be estimated by calculating the means of the sampled values from their posterior probabilities. We also recorded the proportion of iterations that each SNP was assigned to category 2, 3 or 4. If the proportion exceeds the threshold, the SNP is the BayesRB selected associated SNPs.

Then, we made risk prediction of being affected for the individuals in the testing data set. We set 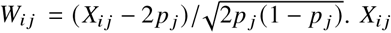 is the number of copies of the risk alleles (0, 1, 2) at SNP *j* for individual *i* with *p_j_* being the frequency of the risk allele in the training population.

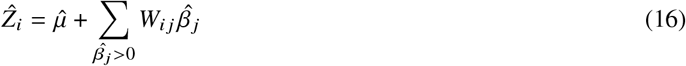

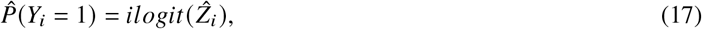

where 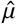 and 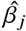 are estimated from the MCMC above. 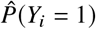 is the estimated probability of being affected for a new individual in the testing data set.

### 2.2 Other Approaches

#### BayesR

BayesR Fortran program [Moser et al., 2015] generates the SNP effect estimates, predicted outcomes, and the proportion of iterations that each SNP is assigned to each category. BayesR SNP effect estimates and the predicted outcomes are directly obtained from the output. The BayesR selected associated SNPs are those with the proportions of iterations that the SNPs are assigned to category 3 or 4 bigger than the threshold (Table 1).

**Table 1:** An overview of how the BayesRB, BayesR, logistic regression and LASSO results are generated.

| Approaches | Criteria to select associated SNPs | Associated SNPs | SNP Effects | Risk Prediction |
| --- | --- | --- | --- | --- |
| **BayesRB** | Proportion of iterations that a SNP is assigned to category 2, 3 or 4 | SNPs with the proportion bigger than the threshold | Generated by bayesRB algorithm | Using all the SNP effect estimates |
| **BayesR** | Proportion of iterations that a SNP is assigned to category 3 or 4 | SNPs with the proportion bigger than the threshold | Generated by bayesR algorithm | Using all the SNP effect estimates |
| **Logistic Regression** | Bonferroni-corrected P values | SNPs have the p values smaller than the threshold | Generated by marginal logistic regression | Among the selected SNPs, using uncorrelated SNPs with smallest p values |
| **LASSO** | SNPs that are not shrunk to 0 |  | None-zero effect SNP estimates are re-estimated by logistic regression | Using all the selected SNPs |

#### Logistic Regression

We used PLINK [Purcell et al., 2007] to conduct all the logistic regression analysis. The SNP effects are estimated by the marginal logistic regression method. The logistic regression selected associated SNPs are those with p values smaller than the Bonferroni-corrected threshold. Among the logistic regression selected associated SNPs, we selected one SNP with the smallest p value in each linkage disequilibrium (LD) block to do the prediction.

#### LASSO

We also used PLINK [Purcell et al., 2007] to conduct all the LASSO analysis. The LASSO selected associated SNPs are those whose SNP effects are not shrunk to zero with the shrinkage parameter *λ*. The SNP effects for the LASSO selected associated SNPs are estimated using marginal logistic regression. The other SNP effects are zero. The predicted disease risks are estimated by the logistic regression using all the LASSO selected associated SNPs.

### 2.3 Study Design

In this study, we proposed a Bayesian approach (BayesRB) to select the associated SNPs, estimate the SNP effects and predict genetic risk for dichotomous traits. We applied the approach to pilot simulated data sets, genome-wide simulated data sets and real data sets. There are two pilot simulated data sets. One has unrealistic SNP effects, while the other has more realistic SNP effects. Using the pilot simulated data sets, we diagnosed the performance of the BayesRB approach. To examine whether the parameters mix well, we assessed the convergence of the parameters. We also explored whether it is necessary to infer the genetic variance from the data and how genetic variance affects the SNP effect estimation. We diagnosed the performance of the BayesRB approach with and without genetic variance fixed. The genome-wide simulated data set is even more like a real data set with some SNPs correlated to each other, showing realistic linkage disequilibrium patterns. Using the genome-wide simulated data set, we aimed to measure BayesRB’s performance when the SNPs are not fully independent of each other. Considering the computational time, we simulated 50 genome-wide simulated data sets. Each has 3000 individuals. For each one, we randomly selected 80% of individuals to the training data set, and the rest to the testing data set, letting equivalent proportions of the cases to the controls in the two data sets. We selected the associated SNPs and estimated the SNP effects using the training data set and predicted the risk of being affected using the testing data set. We used the Crohn’s disease (CD) and bipolar disorder (BD) data set from Welcome Trust Case Control Consortium (WTCCC) version 1 as the real data sets to investigate the BayesRB approach. The WTCCC data sets were outlined in [Consortium et al., 2007]. We created 20 random 80/20 training and testing splits of the two data sets. The purpose of using these two WTCCC data sets is to investigate the performance of BayesRB in real data sets.

The data description and the study design diagram can be found in Figure 3.

**Figure 3:**
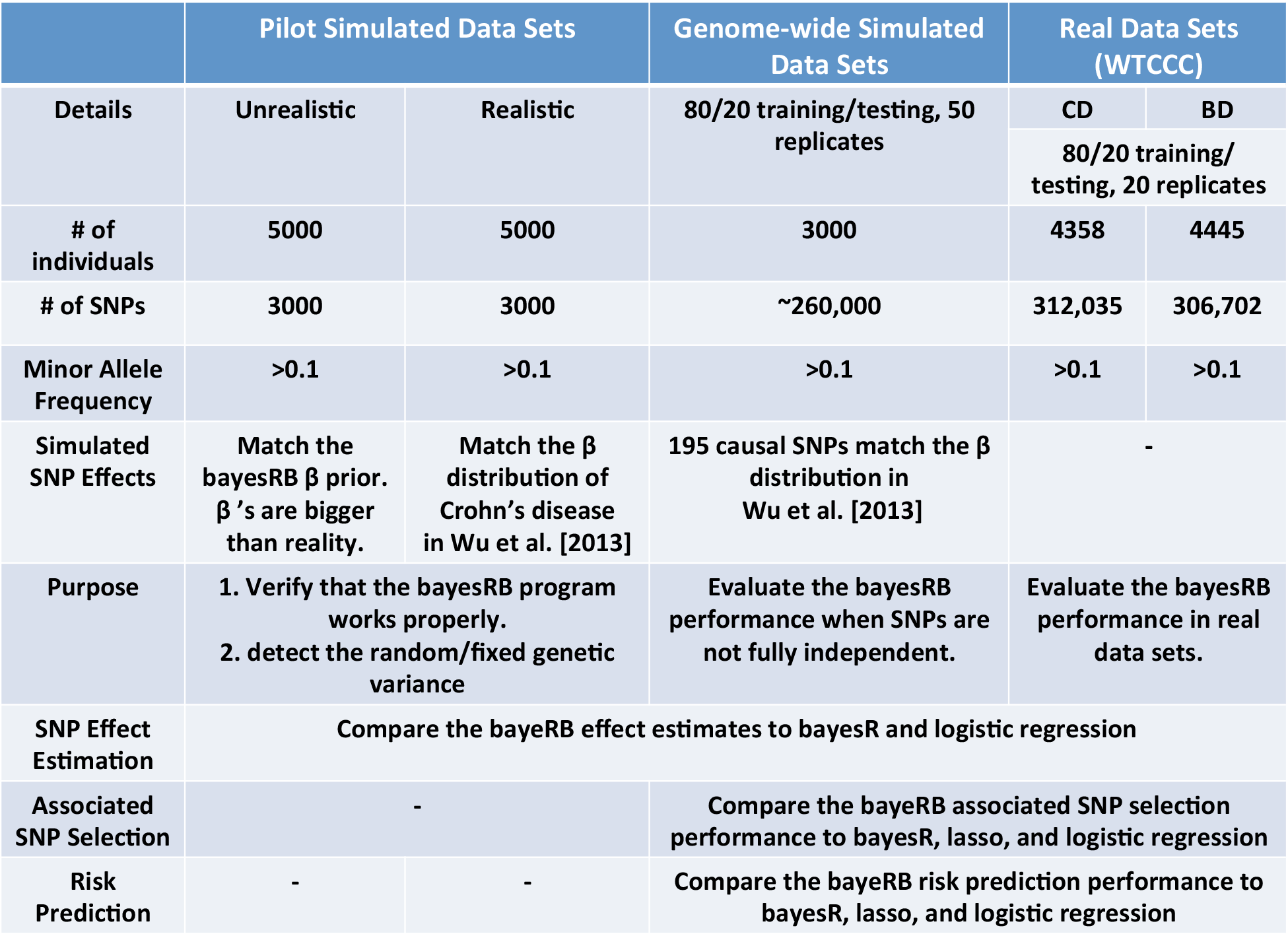
Pilot simulated data sets, genome-wide simulated data sets, and real data sets description and the study design diagram.

## 3 Results

### 3.1 BayesRB R package

We wrote a BayesRB R package using RCPP. All the following results are generated by the BayesRB R package. The source code can be found in the following website: https://github.com/sylviashanboo/BayesRB

### 3.2 Pilot Simulated data sets

To ensure the program works correctly, we first simulated two pilot simulated data sets. In both of the data sets, the SNPs are independent and the individuals are independent.

We simulated both data sets with 5,000 individuals and 3,000 SNPs using the Multiple Gene Risk Prediction Performance (mgrp) R package [Pepe et al., 2010]. One data set has unrealistic SNPs effects. The SNP effects were set bigger than those usually seen in a typical GWAS and matching the prior of ***β*** as Moser et al. [2015] did: 50 SNP effects have variance of 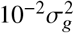, 310 SNP effects have variance of 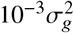, and 2680 SNP effects have variance of 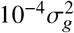, where 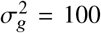. The other data set has more realistic SNPs effects based on the SNP effect distribution of Crohn’s disease indicated in Wu et al. [2013]. We simulated the data set with dichotomous outcome with the proportion of cases to controls of 0.4. The minor allele frequencies were randomly sampled from the abdominal aortic aneurysms (AAA) data set, where AAA data set is a real genotype data of SNPs measured on 3104 individuals. There are 326,706 minor allele frequencies (MAF), which are ≥ 0.1, that can be sampled from.

#### Unrealistic Data

For the unrealistic data set, we ran the the Markov chain for 60,000 cycles with the first 10,000 samples discarded as warm-up. We drew every 50*^th^* sample after warming-up. We first treated 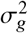 as random. We tuned the variance of the proposal distribution (*θ*) manually. When *θ* = 1, all the parameters mixed well. Then, we treated 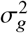 as fixed. We set different values of 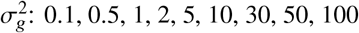, and 500. Figure 4 shows the autocorrelation of the estimated parameters when 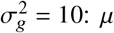, three *β*’s with the biggest absolute value, two randomly selected *λ*’s, and two random selected *Z*’s are all mixed very well. When 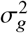 takes the other values, the autocorrelation plots show similar patterns. Figure 5 shows that when 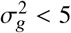, the ***π*** parameters do not mix well and have high autocorrelation. But when 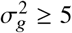, the ***π*** parameters mixed well.

**Figure 4:**
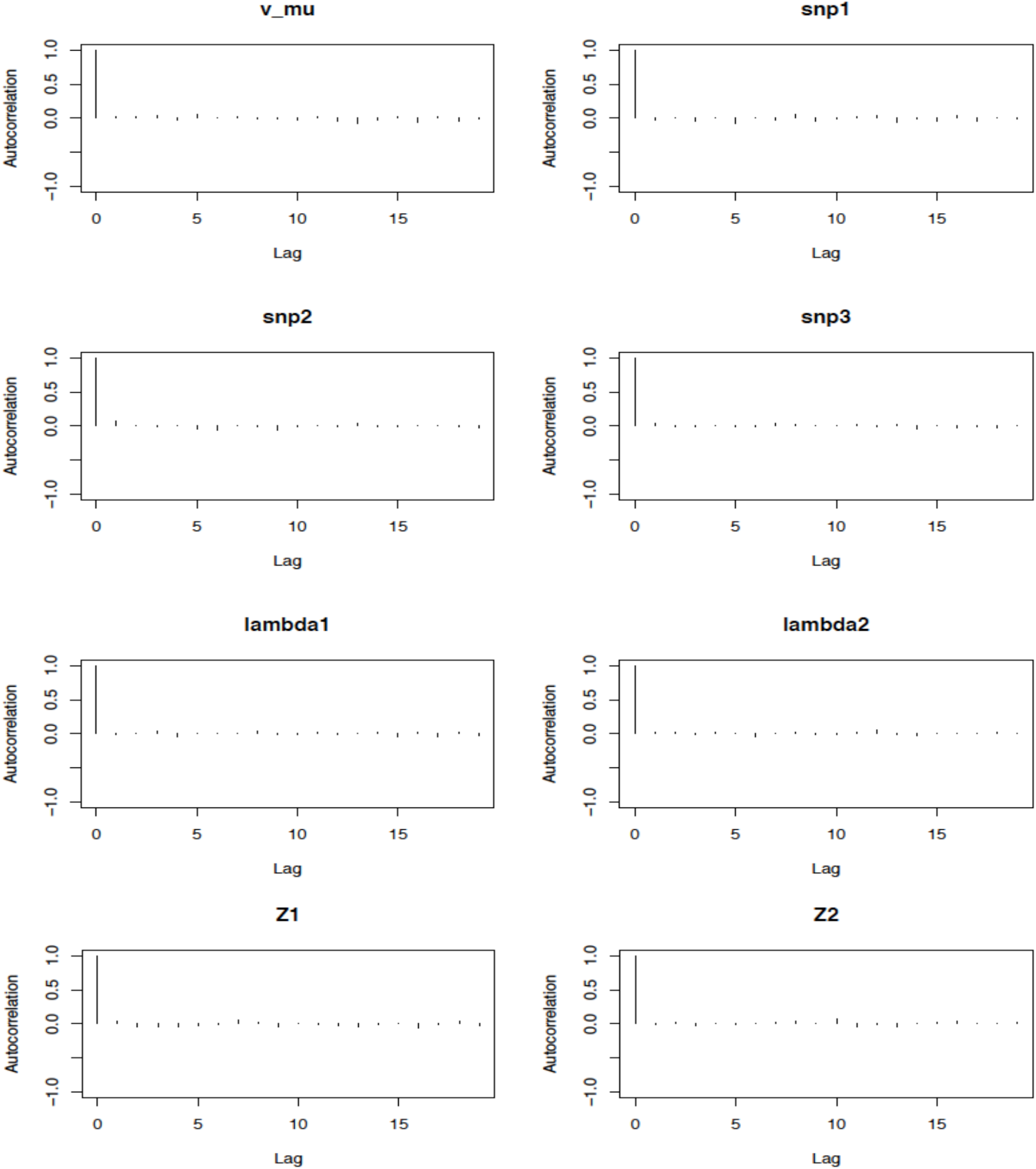
Autocorrelation of the following parameters: *μ*, three *β*’s with the biggest absolute values, two randomly selected *λ*’s, two randomly selected *Z*’s when 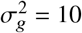.

**Figure 5:**
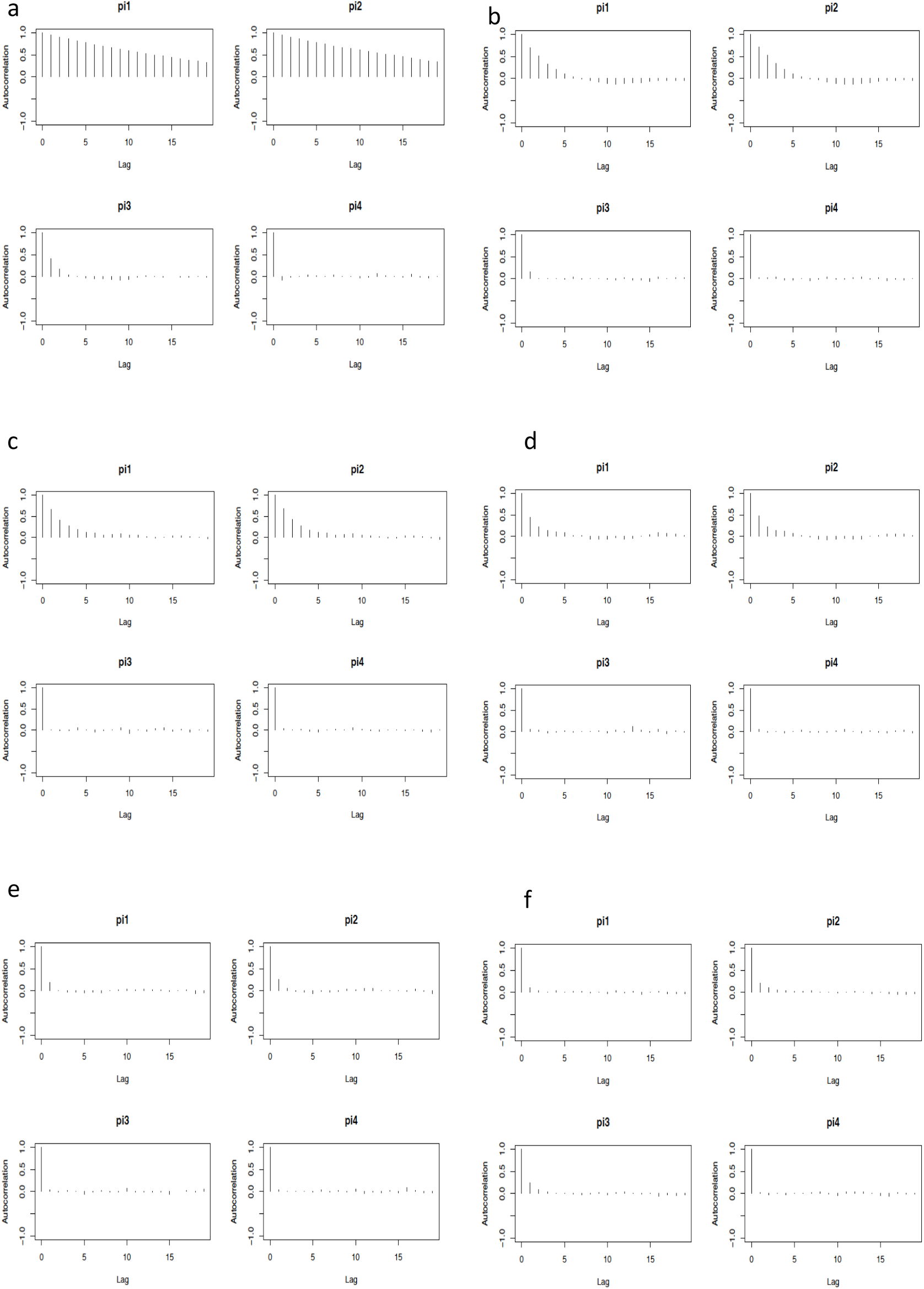
Autocorrelation of *π* when 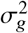 is 0.1 (a), 0.2 (b), 0.5 (c), 1 (d), 5 (e), and 10 (f).

We compared the BayesRB ***β*** estimates to the logistic regression ***β*** estimates, which are considered unbiased. Figure 6 shows that when 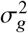 is set to a fixed value smaller than or equal to 10, the ***β*** estimates are under estimated, compared to the logistic regression estimates. When 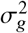 is set to a fixed value bigger than 10 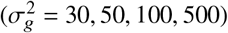 or is estimated using Metropolis-Hasting sampling, their ***β*** estimates of a given SNP are similar. In this situation, BayesRB shrunk the small SNP effects to zero, and estimated the bigger SNP effects similar to the logistic regression estimated ***β***.

**Figure 6:**
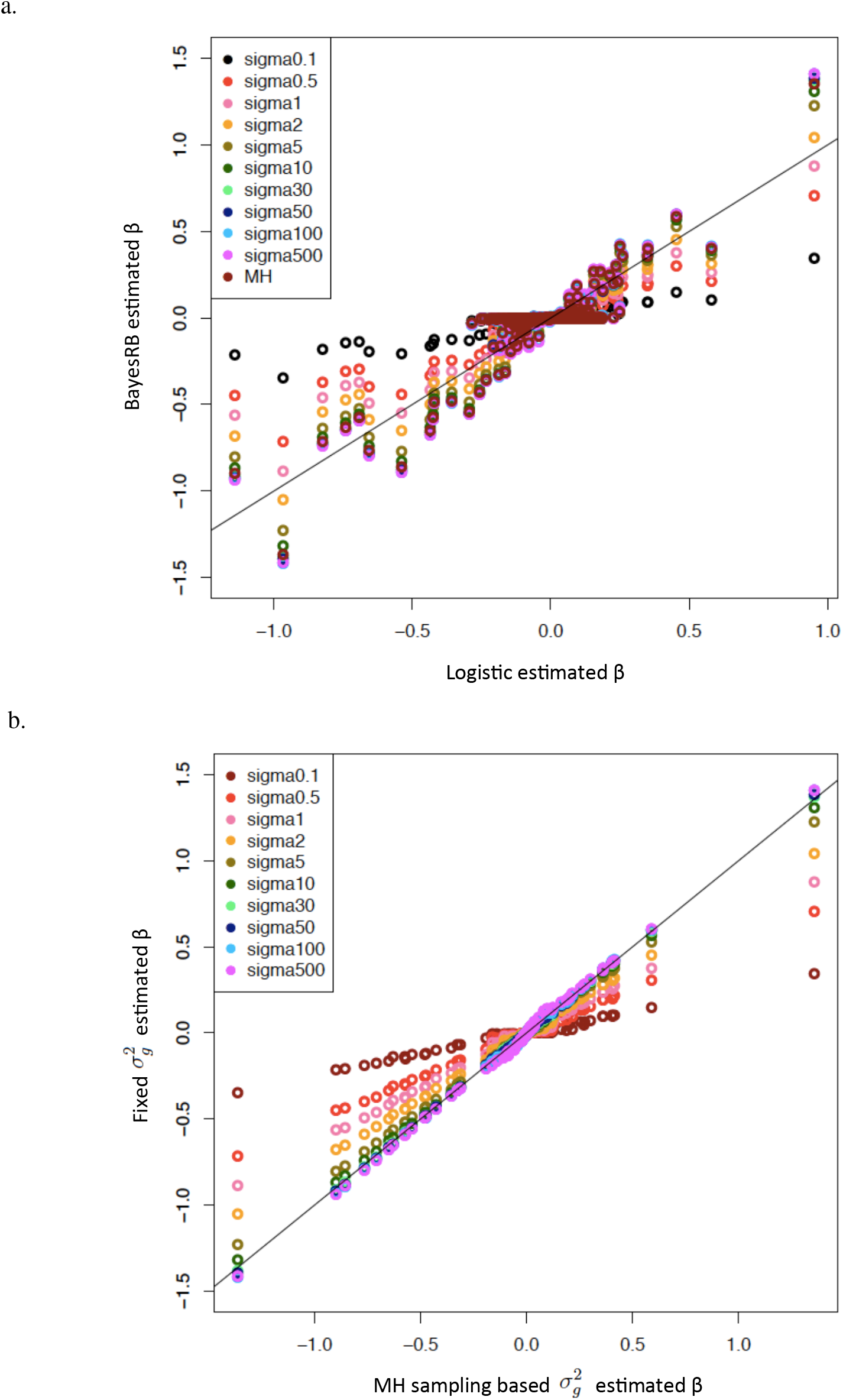
A comparison of the SNP effect estimates under different 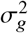 using the unrealistic data set. **a.** A comparison of the BayesRB *β* estimates to the logistic regression *β* estimates. **b.** A comparison of the *β* estimates with 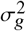 updated by Metropolis-Hasting sampling to the *β* estimates with 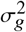 fixed at different values. The solid lines in both plots are the diagonal lines.

From the recorded category that each SNP is assigned to, in each MCMC loop, we found that the relatively big effect SNPs are more often assigned to the category 2, 3 or 4. Figure 7 shows that when 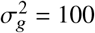, more than 90% of the SNPs with the logistic regression estimated ***β*** bigger than 0.25 are assigned more than 95% of the time to category 2, 3 or 4 by BayesRB.

**Figure 7:**
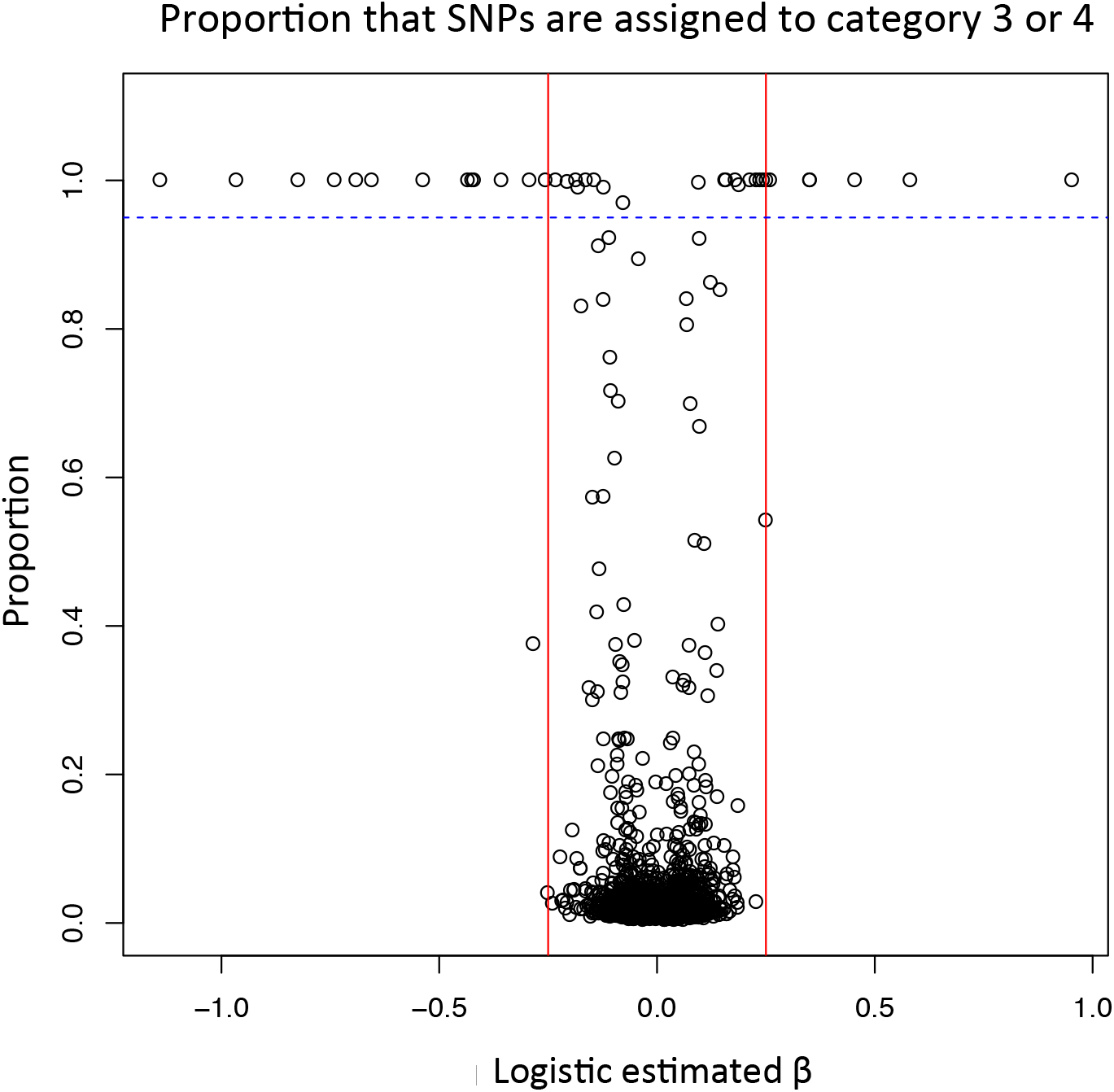
The proportion of iterations that each SNP is assigned to category 2, 3 or 4, when 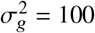. The blue dotted horizontal line indicates the proportion of 95%. The red vertical lines are drawn at −0.25 and 0.25.

#### Realistic Data

For the realistic data set, we ran the the Markov chain for 20,000 cycles with the first 10,000 samples discarded as warm-up. We drew every 10*^th^* sample after warming-up. We treated 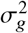 as fixed. The autocorrelation plots show similar patterns as that using unrealistic data. When 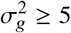, all the parameters including ***π*** mixed well.

We also compared the BayesRB ***β*** estimates to the logistic regression ***β*** estimates. For the SNPs with the absolute value of the logistic regression ***β*** estimates smaller than 0.1, their BayesRB ***β*** estimates are shrunk to a value close to 0 (Figure 8a). For the SNPs with the absolute value of the logistic regression ***β*** estimates bigger than 0.1, the small 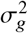 values 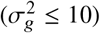 will let the ***β*** underestimated, while big 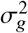 values (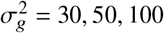, or 500) have ***β*** estimates similar to the logistic regression ***β*** estimates. (Figure 8c, d).

**Figure 8:**
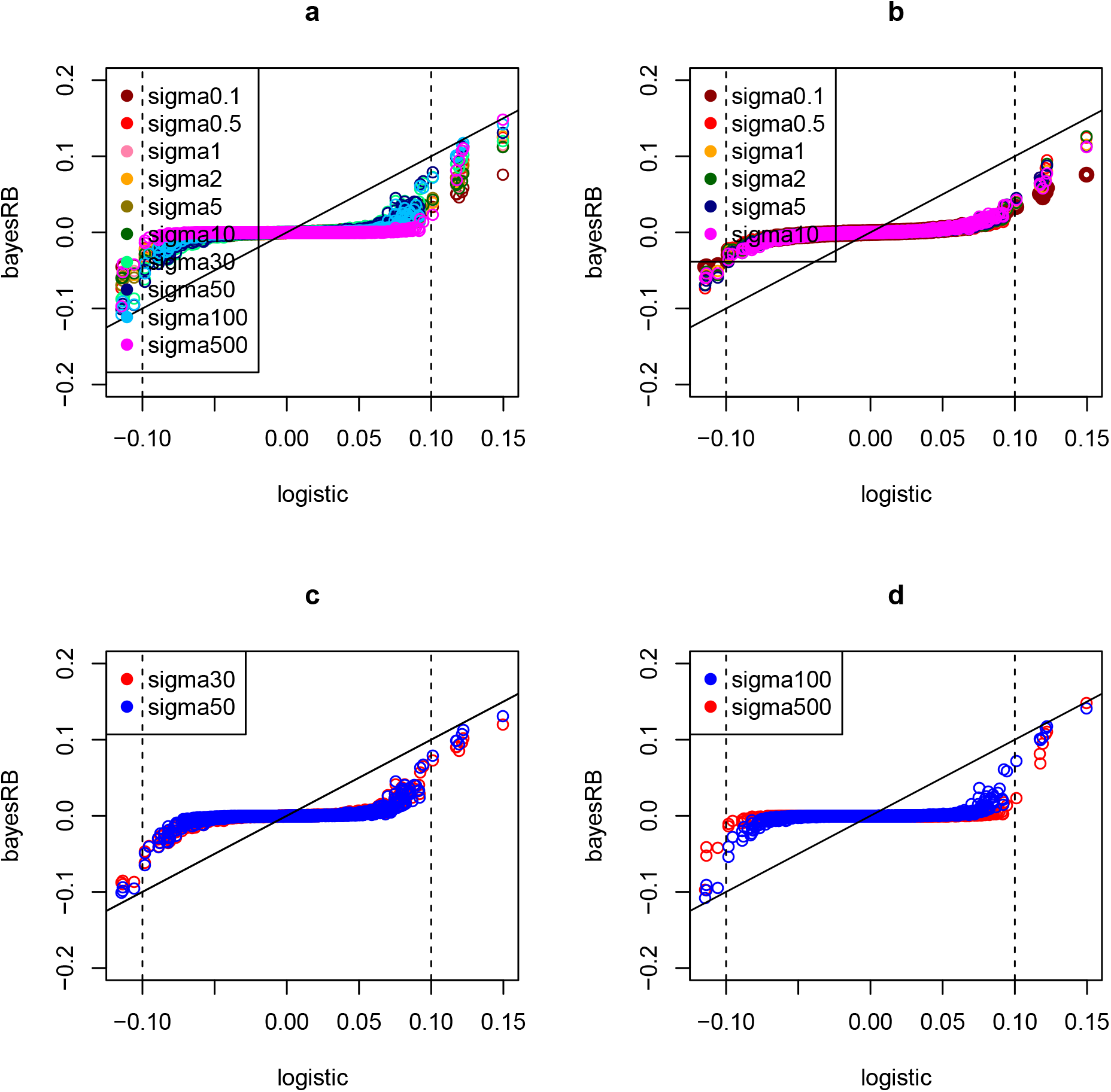
A comparison of the SNP effect estimates under different 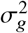 using the realistic data set. A comparison of the BayesRB *β* estimates to the logistic regression *β* estimates. The x axis label “logistic” indicates the SNP effects estimated by logistic regression. The y axis label “BayesRB” indicates the SNP effects estimated by BayesRB. The solid line in each plot is the diagonal line showing the equivalent values of logistic regression estimated *β* and the BayesRB estimated *β*. The dotted lines indicate the logistic regression estimated *β* of 0.1 and −0.1, respectively.

When the 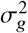 is large enough 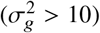, different values of 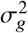 affect the shrinkage level of the ***β*** estimates. Figure 9a shows that, for the SNPs with the logistic regression ***β*** estimates smaller than 0.1, the bigger the 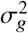, the smaller the summation of the BayesRB ***β*** estimates squared. In other words, for the SNPs with the logistic regression ***β*** estimates smaller than 0.1, when 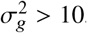, the bigger the 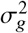, the more the BayesRB ***β*** estimates are shrunk. Figures 9b shows that, for the SNPs with the logistic regression ***β*** estimates smaller than 0.1, when 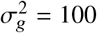, the overall distance of BayesRB ***β*** estimates and logistic ***β*** estimates is the smallest. In other words, when 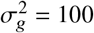, the BayesRB ***β*** estimates for the big effect SNPs are closest to the logistic ***β*** estimates. Therefore, we set the 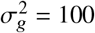 for the BayesRB analysis in genome-wide simulated data set and the real data set.

**Figure 9:**
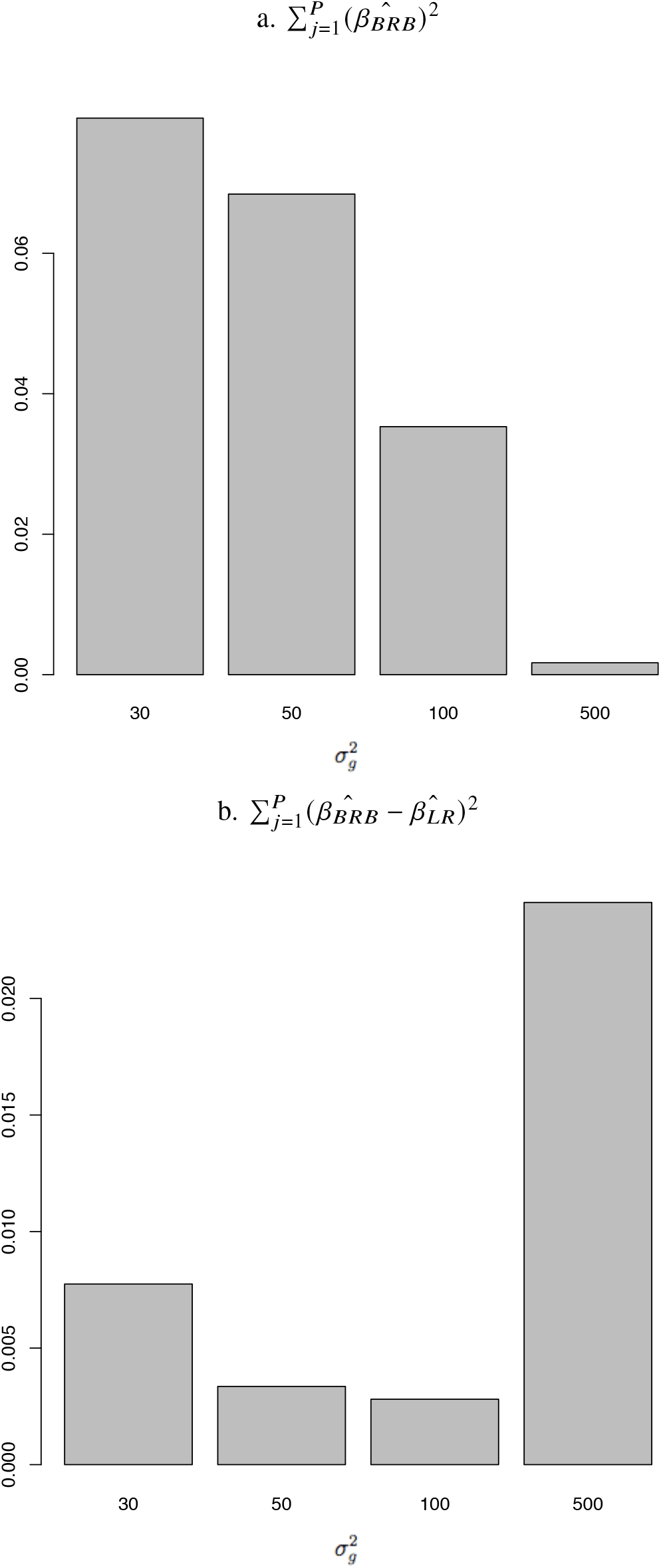
**a.** 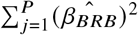, when 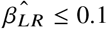. **b.** 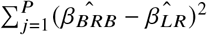, when 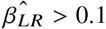. In the formula, 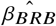 is the BayesRB estimated SNP effect; and 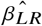 is the logistic regression estimated SNP effect.

### 3.3 Genome-wide Simulated data sets

To measure how well BayesRB works on genome-wide data sets with SNPs that are not fully independent, we applied BayesRB to genome-wide simulated data sets. We first simulated a population genetic data set containing 16,000 individuals’ genotype data set using the software GWAsimulator [Li and Li, 2008] based on the HapMap phased CEU data. The genotype data contains all the 305,054 SNPs in the Illumina HumanHap300 SNP chip. We filtered out the SNPs in the data set with MAF smaller than 0.1. We assigned 195 causal SNPs that are not in the same LD blocks in the genome with effects follow a distribution described in Wu et al. [Wu et al., 2013]. The top 6 biggest effect causal SNPs (absolute values of the assumed true SNP effects ≥ 0.1) are defined as big effect causal SNPs. The 7 to 21 biggest effect causal SNPs (0.05 ≤ absolute values of the assumed true SNP effects < 0.1) are defined as the medium+ effect causal SNPs. The 22 to 34 biggest effect causal SNPs (0.038 ≤ absolute values of the assumed true SNP effects < 0.05) are defined as the medium-effect causal SNPs. Medium+ effect causal SNPs combined with medium-effect causal SNPs are medium effect causal SNPs. The rest SNPs (absolute values of the assumed true SNP effects < 0.038) are small effect causal SNPs. We obtained 1,500 cases and 1,500 controls from all the 16,000 individuals using the software GCTA [Yang et al., 2011], setting the heritability equal to 0.5 and disease prevalence equal to 0.1. We split each data set to 80/20 training and testing data sets, each with the same proportions of cases and controls.

First, we explored how close the genome-wide simulated data sets are to the real genome-wide data sets. Figure 10 is based on the logistic regression result of one of the genome-wide simulated data sets. In Figure 10a, the SNP effect density is close to real studies shown in Wu et al. [2013]. Figure 10b and c show that the big effect causal SNPs have smallest p values and the biggest SNP effect estimates. And there are peak towers around the big effect SNPs due to the LD, which is similar to a real GWAS manhattan plot. Therefore, the genome-wide simulated data sets are similar to the real genome-wide data sets.

**Figure 10:**
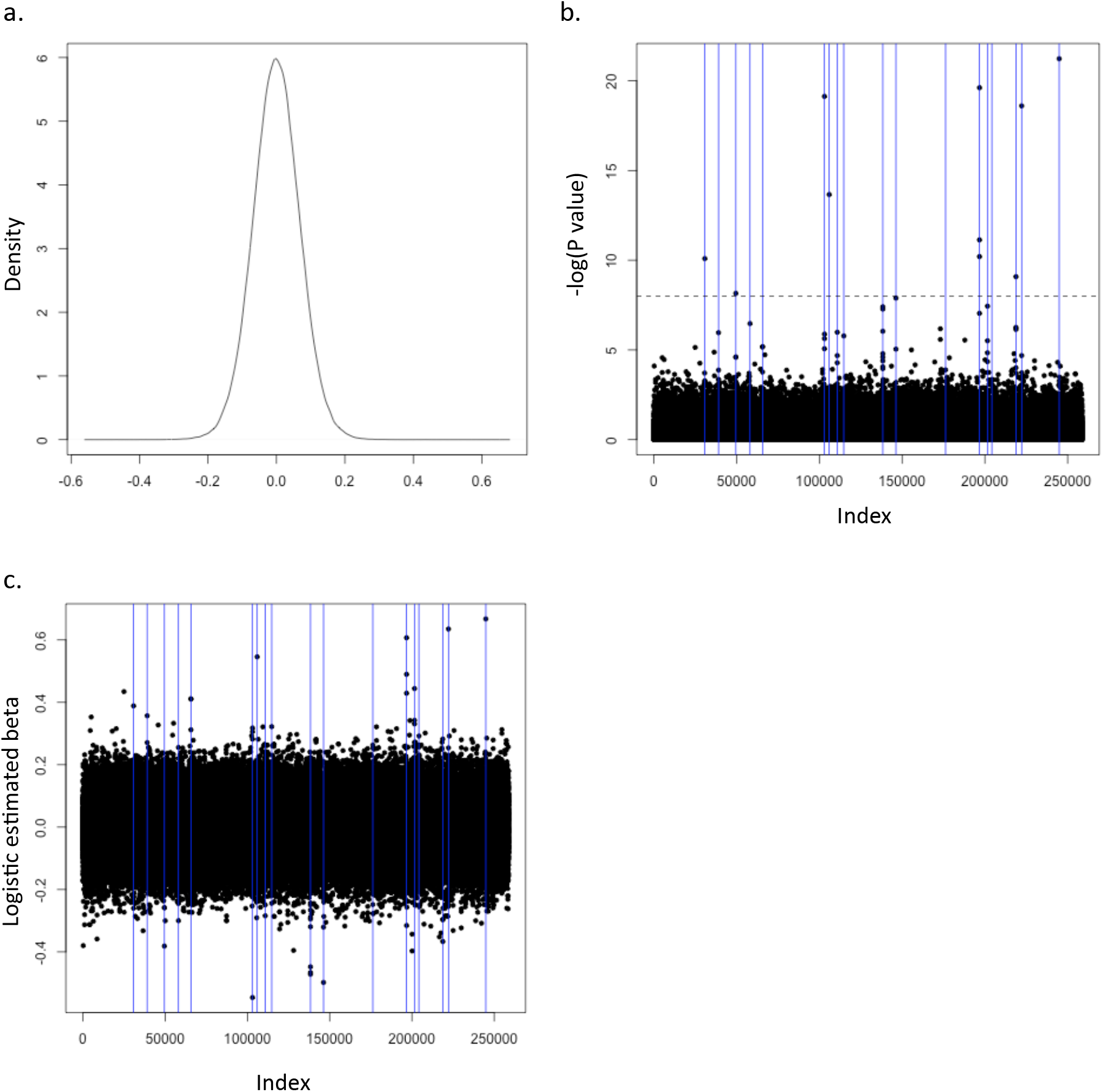
Evaluation of the genome-wide simulated data sets by visualizing the distribution of SNP effect estimates and −*log*_10_(P values) generated by logistic regression method. **a.** Density plot of the logistic regression SNP effect estimates. **b.** Scatter plot of −*log*_10_(P values) generated by logistic regression method. The vertical blue lines indicate the top 21 SNPs with the biggest causal SNP effects (big effect causal SNPs and the medium+ effect causal SNPs). The horizontal dotted line indicates −*log*_10_ (*P*) = 5 × 10^8^. **c.** Scatter plot of SNP effect estimates by logistic regression method. The vertical blue lines have the same meaning as b.

For each BayesRB analysis on genome-wide simulated data sets, we ran the Markov chain for 15,000 cycles with the first 5,000 cycles as warm-up. We drew every 20 sample after warming-up. We made diagnostic plots. The autocorrelation plots of most parameters show similar patterns as Figure 4, which indicates that the parameters are mixed well. But some do not. A larger number of MCMC loops and the bigger thinned number is required. But considering the computational time, we still used those estimates in the following analysis, although they are not ideal to use.

We randomly selected one data set out of 50 and compared the SNP effect estimation performance of BayesRB to BayesR, logistic regression and LASSO method using this data set. Since LASSO underestimates the selected associated SNP effects, we reestimated the SNP effects by using the marginal logistic regression analysis. Then, LASSO and logistic regression methods provide the same SNP effect estimates. So, we only showed logistic regression result here. Figure 11a shows that BayesRB estimates have a linear relationship with the BayesR estimates, but BayesR has shrunk estimates. Figure 11b shows that the SNPs with big logistic regression estimates have BayesRB estimates close to logistic regression estimates. SNPs with small logistic regression estimates have BayesRB estimates close to 0. The red dots, which indicate the big effect and the medium+ effect causal SNPs, show that the top 21 biggest effect causal SNPs have relatively big effect estimates. Figure 12 shows the proportion of the loops that the SNPs are assigned to the category 2, 3 or 4. All the big effect and the medium+ effect causal SNPs have relatively big proportions, which are bigger than 0.2.

**Figure 11:**
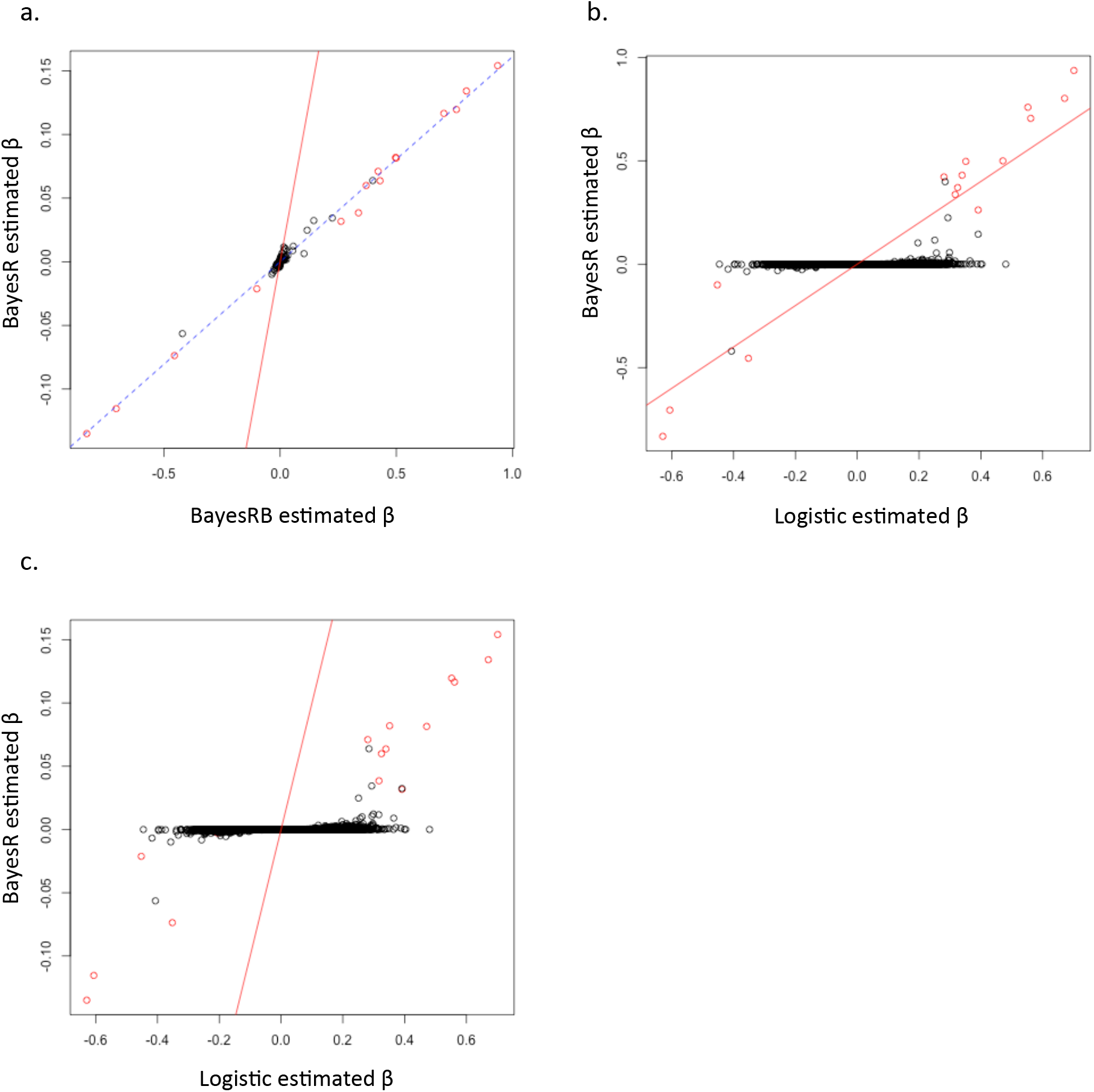
The comparisons of SNP effect estimates by BayesRB, BayesR, and logistic regression detected in one of the genome-wide simulated training data sets. The red dots indicate the SNP effect estimates of the 21 biggest effect SNPs (big effect causal SNPs and the medium+ effect causal SNPs). The red solid line is the diagonal line. **a.** BayesRB SNP effect estimates vs. BayesR SNP effect estimates. The blue dotted line is the fitted regression line. **b.** Logistic regression SNP effect estimates vs. BayesRB SNP effect estimates. **c.** Logistic regression SNP effect estimates vs. BayesR SNP effect estimates.

**Figure 12:**
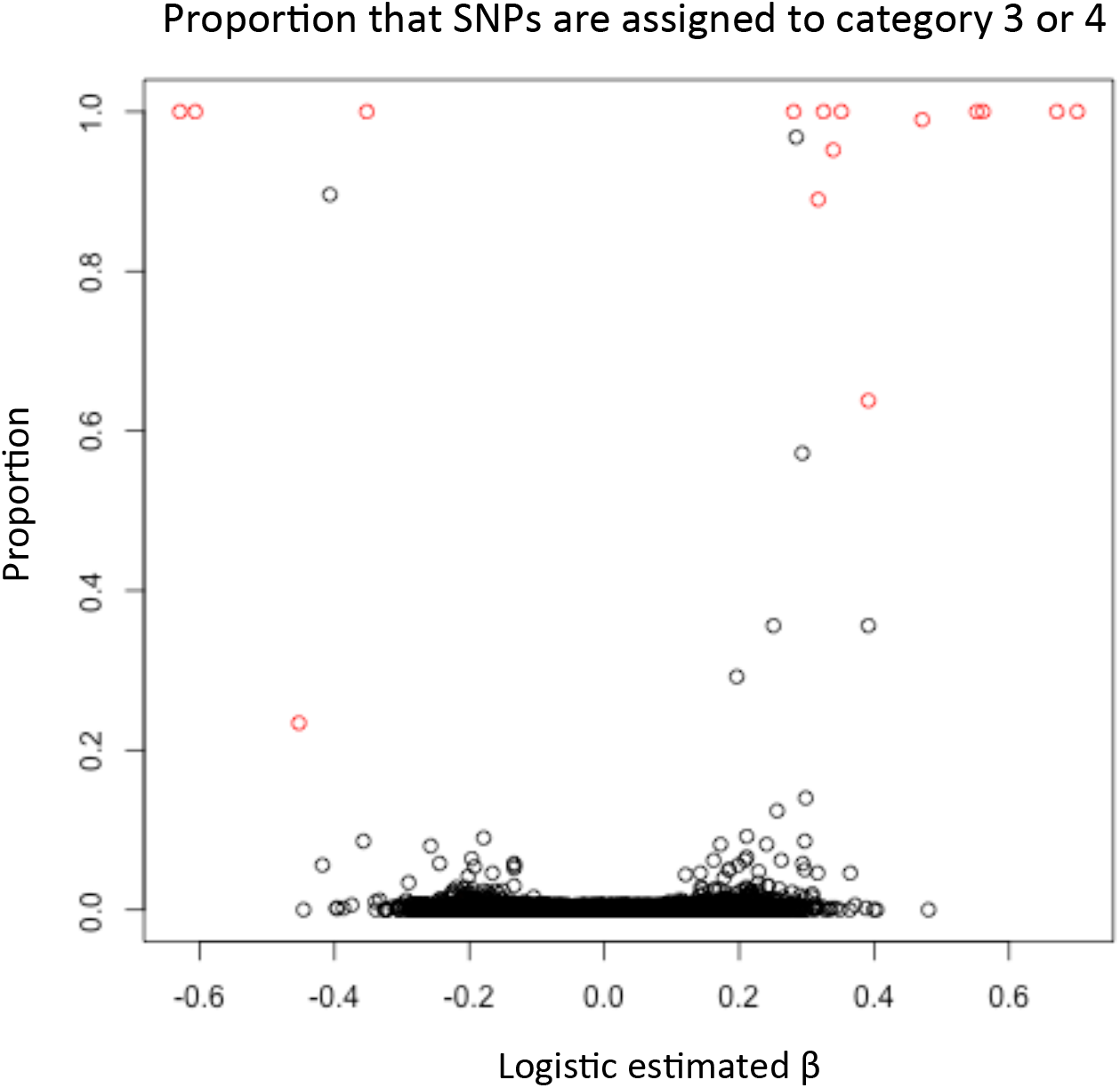
The proportion of the iterations that the SNPs are included in the model in one of the genome-wide simulated training data sets. The red dots indicate the 21 biggest effect SNPs (big effect causal SNPs and the medium+ effect causal SNPs).

We also compared the performance of identifying associated SNPs of BayesR, BayesRB, logistic regression and LASSO. Comparison between methods are assessed on their ability to identity genomic region of 50 SNPs window containing causal SNPs. The reason we used 50 SNPs window is because we did not have the SNP manifest identifying the SNP locations for the Illumina HumanHap300 phased data set we used in the GWASimulator software. And on the Illumina website, they indicated “Although assays on the Human Hap300 BeadChip were chosen using tagSNPs, SNPs are evenly spaced across the genome to ensure comprehensive coverage. On average, there is 1 SNP every 9 kb across the genome (median spacing = 5kb). The average 90th percentile gap on the HumanHap300 BeadChip is 19kb.” Therefore, it is reasonable to set the window size as that containing 50 SNPs. We calculated the true positive rate (TPR) and the false positive rate (FPR) of the four methods to detect the windows containing causal SNPs in each replicate. The TPR is the proportion of windows containing causal SNPs that are correctly identified as containing associated SNPs. The FPR is the proportion of windows not containing causal SNPs but are incorrectly identified as containing associated SNPs. Figure 13 shows that all the methods have mean TPR of 1 to detect the windows containing the five big effect causal SNPs. (One big effect causal SNP is excluded in the QC process, thus is not shown in the result.) When the FPR < 0.015, BayesR has the biggest mean TPR to detect the windows containing medium effect causal SNPs. When the FPR >= 0.015, LASSO has the biggest mean TPR. BayesRB has the lower, but not much lower, mean TPR compared to the other methods under the same FPR.

**Figure 13:**
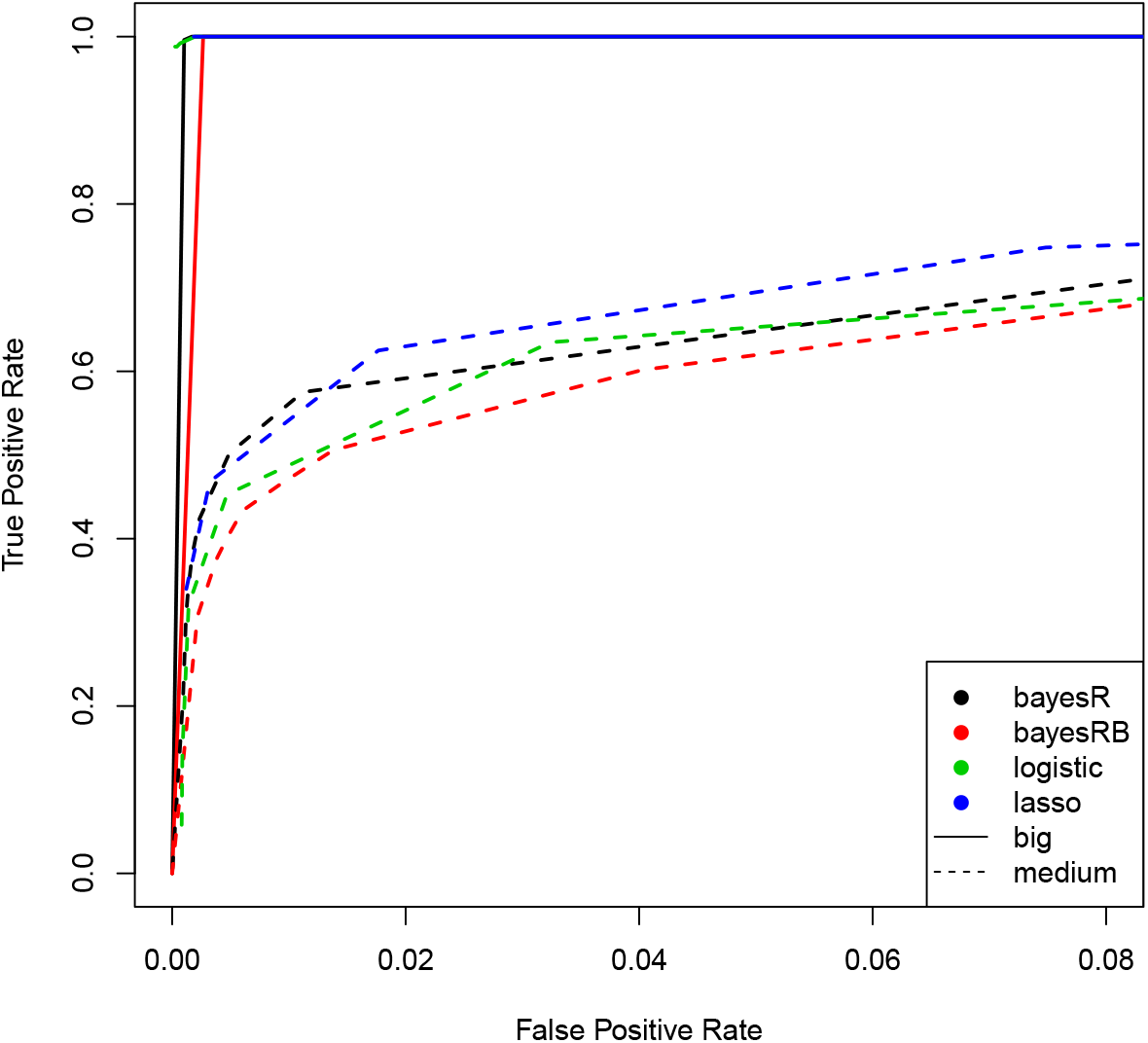
True positive rate vs. false positive rate to detect the windows containing the big effect causal SNPs and the medium effect causal SNPs in the genome-wide simulated data sets.

Then, we compared the power of BayesR, BayesRB, logistic regression and LASSO of having each window containing causal SNPs identified within the 50 replicates. The power is calculated using the number of replicates in which a window containing a given SNP is identified divided by the total number of replicates where at least one SNP in the window have genotype data. Figure 14a and Figure 15a, b, c, and d show that using the thresholds under the same FPR of 0.001, the power of the four methods are similar. LASSO has slightly bigger power for the medium+ effect causal SNPs than the others. BayesRB has slightly bigger power for the medium-effect causal SNPs and the small effect causal SNPs than the others. Figure 14b and Figure 15e, f, g, and h show that using the thresholds under the same FPR of 0.05, LASSO has the biggest power, followed by BayesRB, while BayesR has the lowest power for all the causal SNPs. The power of all the methods detecting the big effect SNPs are 1.

**Figure 14:**
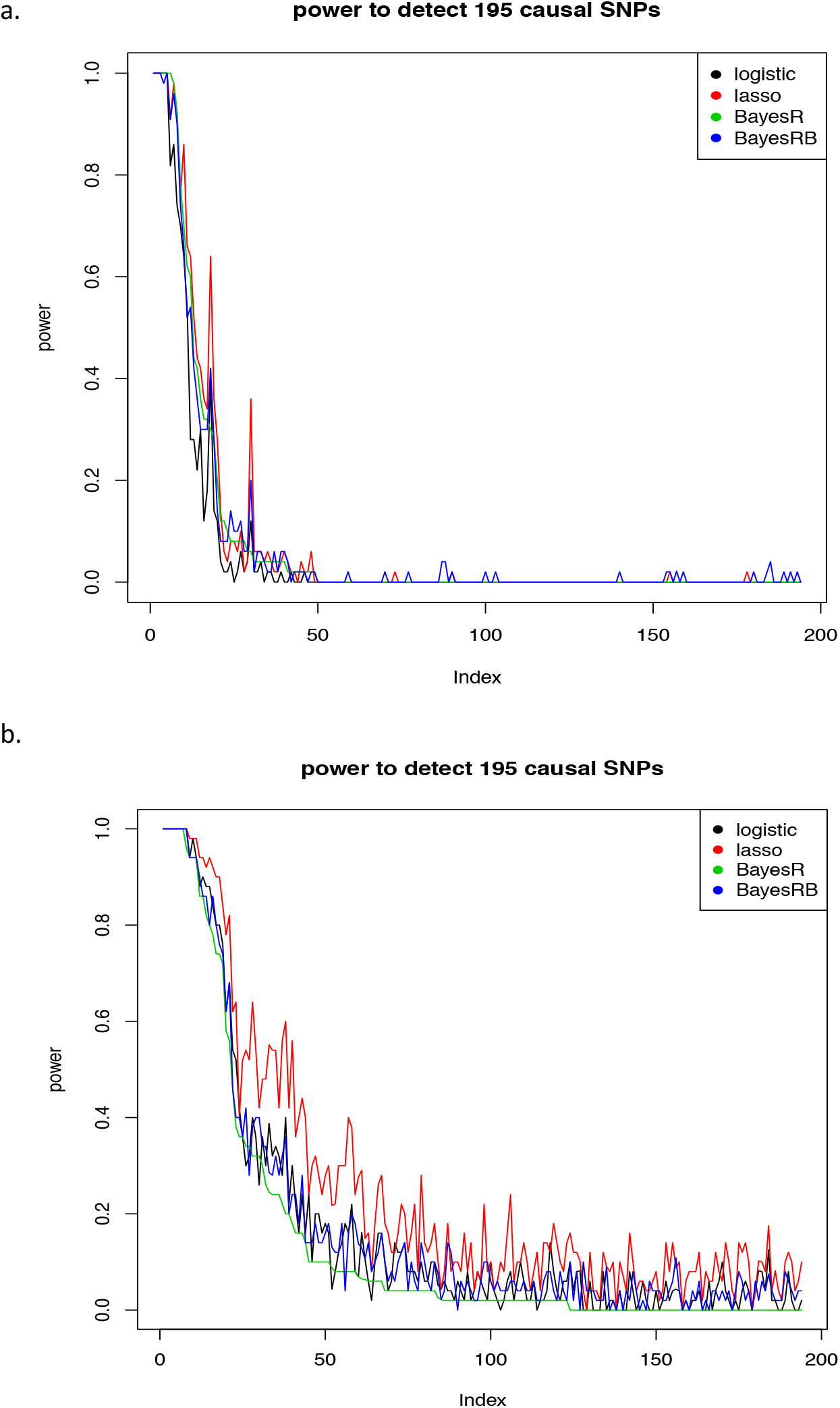
Power of the windows containing the causal SNPs being detected within 50 replicates in the genomewide simulated data sets under the FPR of 0.001 (a) and 0.05 (b). The x axis is the SNP index ordered by the causal SNP effect sizes.

**Figure 15:**
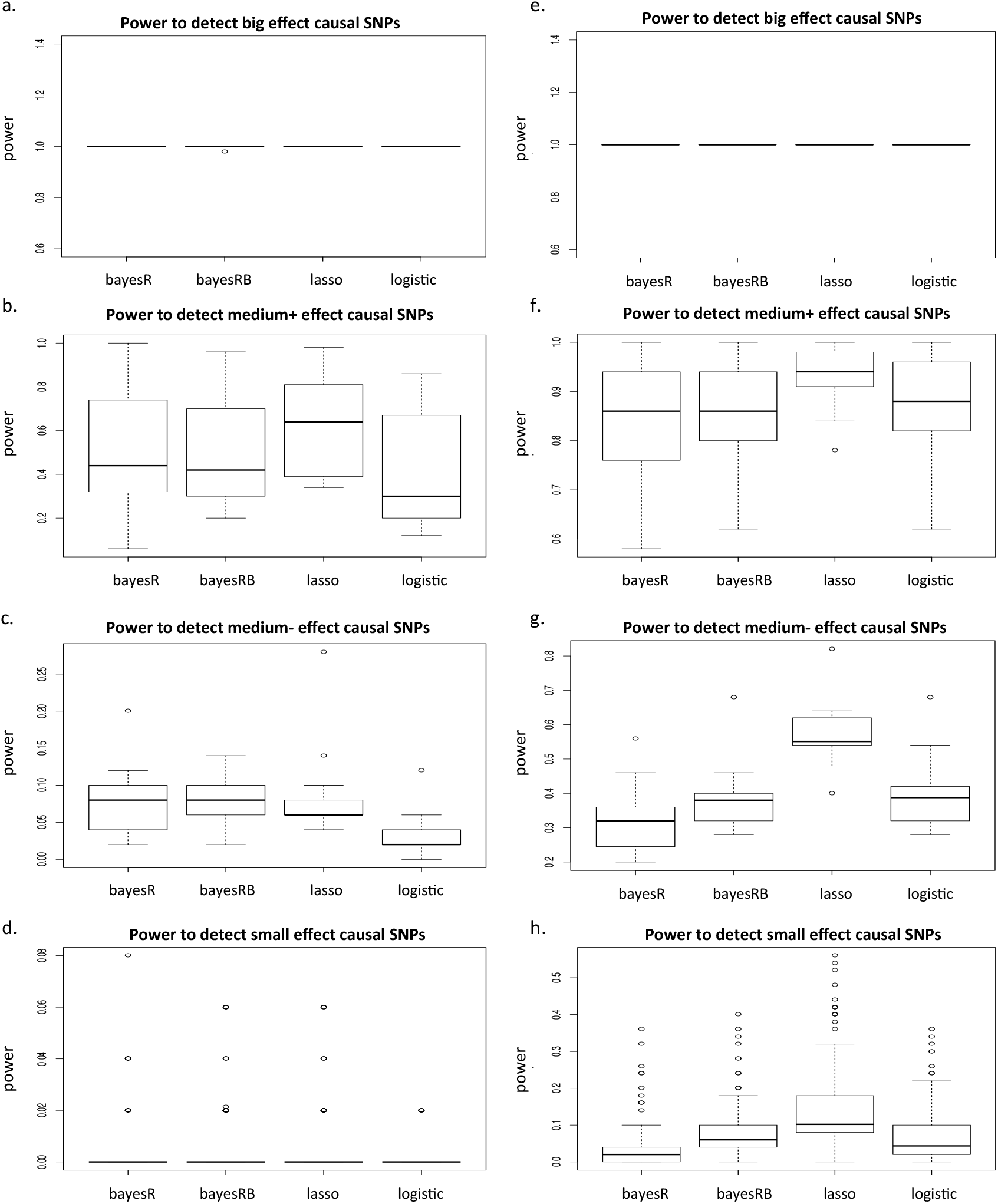
Power of the windows containing big effect causal SNPs (a, e), medium+ effect causal SNPs (b, f), medium-effect causal SNPs (c, g), and small effect causal SNPs (d, h) being detected within 50 replicates under the FPR of 0.001 (a, b, c, d) and 0.05 (e, f, g, h) in the genome-wide simulated data sets.

At last, we compared the performance of risk prediction of BayesR, BayesRB, logistic regression and LASSO on the testing data sets by comparing the area under the curves (AUCs). Figure 16 shows that BayesRB and BayesR generate bigger AUCs than LASSO and logistic regression. But BayesR’s median value of AUC is slightly higher than BayesRB and BayesR’s prediction is more precise than BayesRB.

**Figure 16:**
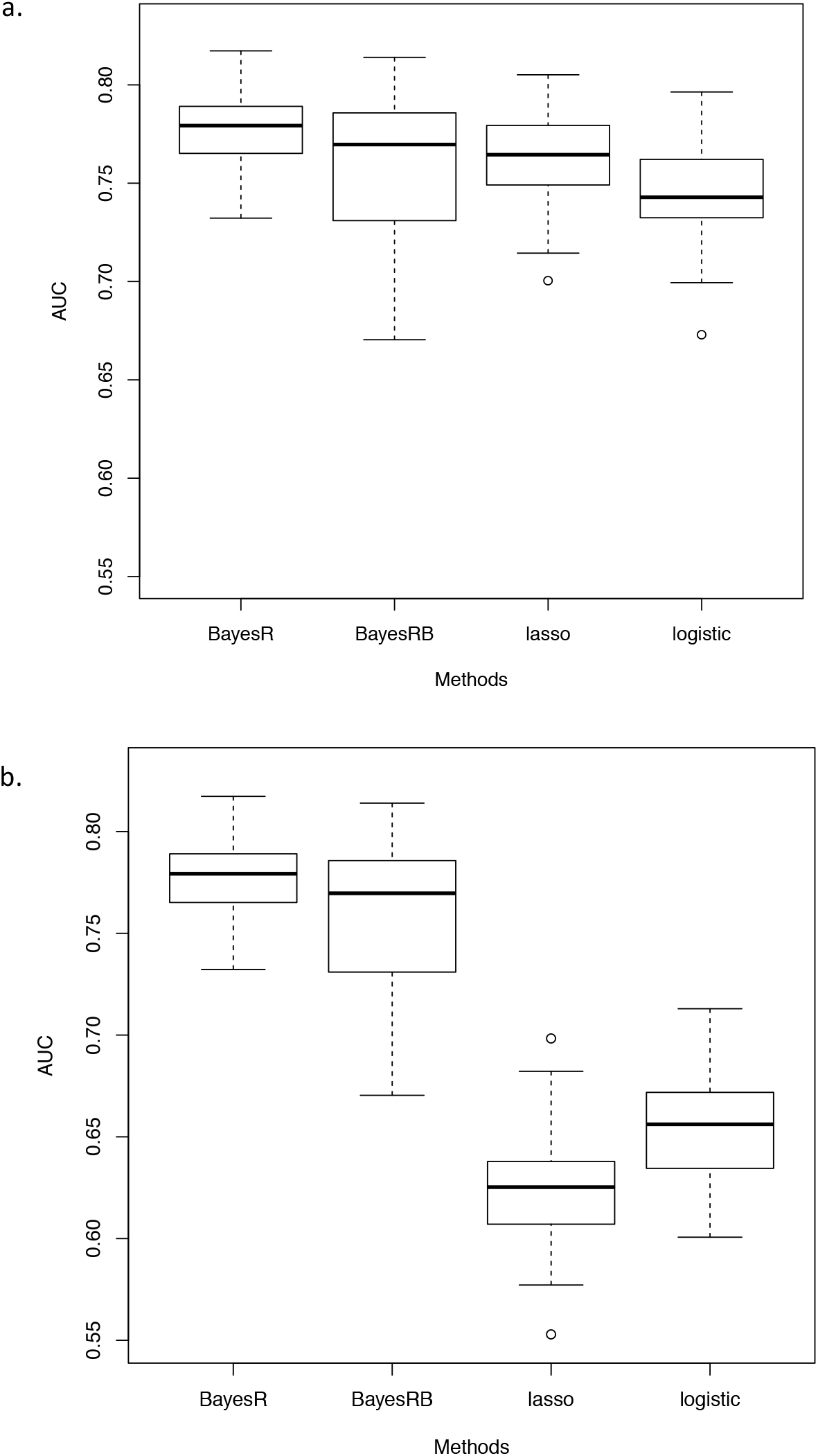
Area under the curve (AUC) of 50 replicates in the genome-wide simulated data sets. Logistic regression and LASSO use the thresholds when FPR = 0.001 (a) and FPR = 0.05 (b).

### 3.4 WTCCC data sets

We assessed the performance of BayesRB for seven diseases of the WTCCC data sets. First, we converted the data sets to PLINK format using MEGA2 [Mukhopadhyay et al., 2005]. Then, following the analyses of Moser et al., we performed strict quality control (QC) on SNP data using PLINK [Purcell et al., 2007]. We removed individuals with missing genotypes greater than 2%. We removed loci with the minor allele frequencies smaller than 1% and SNPs with missingness bigger than 1% for each of the 7 case data sets and the two control data sets. We combined each case and the two control sets into 7 trait case-control studies. We removed the SNPs significant at 5% for differential missingness between cases and controls and SNPs significant at 5% for Hardy-Weinberg equilibrium. We also took relatedness testing using a pruned set of SNPs with LD of *r*^2^ smaller than 0.05. We used the software PRIMUS [Staples et al., 2014] to identify a maximum unrelated set of individuals. We conducted principle component analysis (PCA) using the software GCTA [Yang et al., 2011] and removed individuals in each disease who had poor clustering by visual inspection. Then, we used the software BEAGLE [Browning and Browning, 2016] to fill in the missing genotypes. After QC, the data included 1665 cases of bipolar disorder (BD), 1882 cases of coronary artery disease (CAD), 1576 cases of Crohn’s disease (CD), 1805 cases of hypertension (HT), 1721 cases of rheumatoid arthritis (RA), 1850 cases of type 1 diabetes (T1D), 1761 cases of type 2 diabetes (T2D), and 2757 to 2782 controls depending on the traits. The number of genotypes ranged from 306,702 for BD to 312,035 for CD.

Moser et al. [2015] applied BayesR to all the above seven case-control data sets, treating the binary outcome as the response in an ordinary linear regression. We only used the CD and BD data sets. We chose CD and BD data sets for the following two reasons: 1) both diseases have several significant SNPs contributing to the diseases together, unlike RA or T1D, who are largely influenced by MHC region [Burton et al., 2007]; 2) in Moser et al., BayesR has relatively better performance on these two data sets than the others, except RA and T1D data sets. For both CD and BD data sets, we split each one to 80/20 training and testing data sets for 20 replicates with the same proportions of cases and controls.

Before applying BayesRB to the data sets, we diagnosed the two cleaned data sets to double check the quality control (QC) results. We plotted the Manhattan plots (Figure 17a,c) and the QQ plots (Figure 17b,d) for the two data sets, and compared the plots as well as the significant SNPs to the previous studies. The Manhattan plots and QQ plots, as well as the significant SNPs match the results in previous studies [Burton et al., 2007; Bowden and Dudbridge, 2009]. Therefore, the two data sets are good to use.

**Figure 17:**
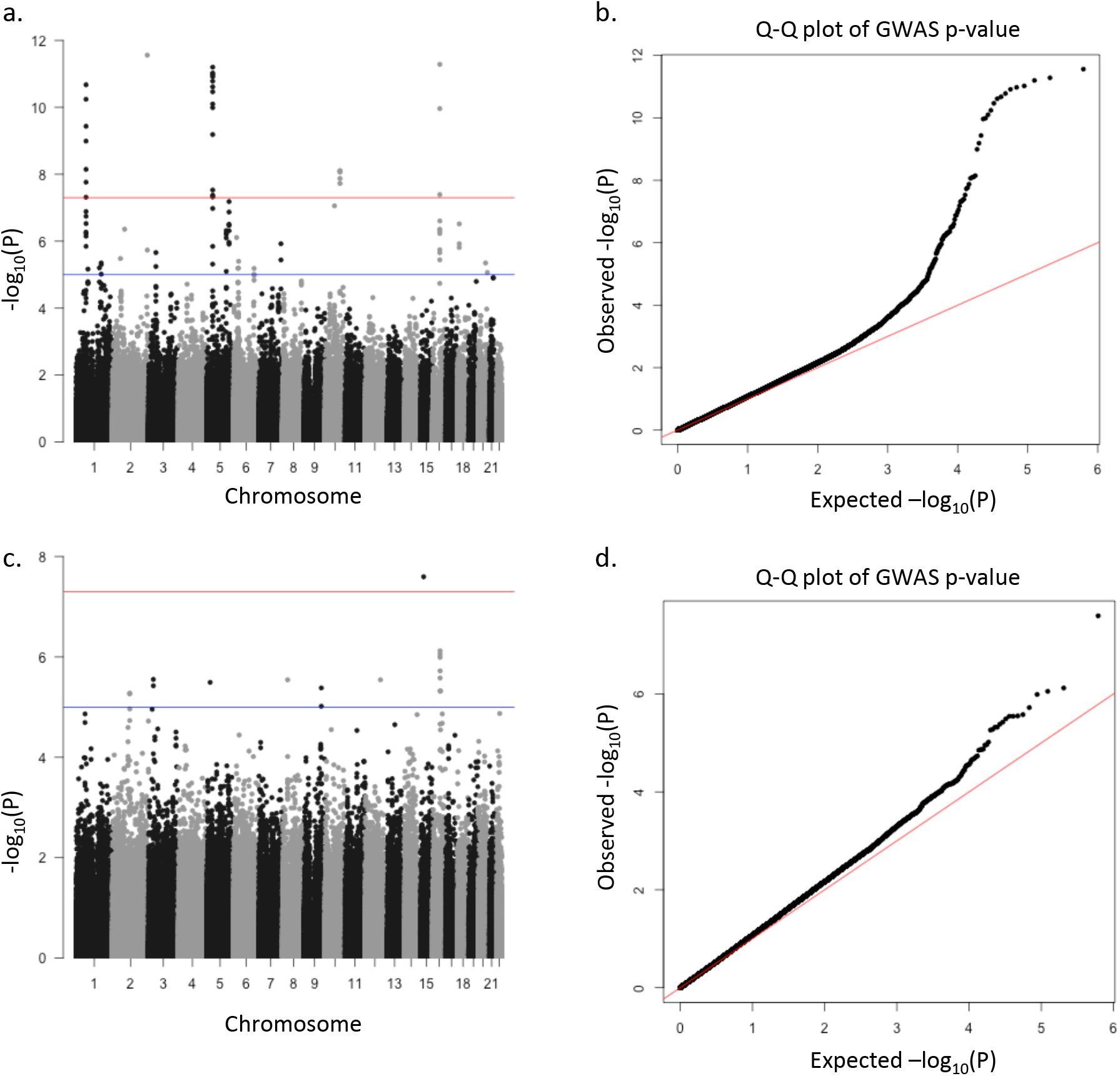
Manhattan plots (a, c) and the QQ plots (b, d) for the CD data set (a, b) and the bipolar disorder data set (c, d).

Both CD and BD data sets have around 45% more individuals and 20% more SNPs than the genome-wide simulated data sets have. Therefore, considering the computational time, we ran the Markov chain for 13,000 cycles with the first 5,000 cycles as warm-up. We drew every 20 sample after warming-up. Then, we made diagnostic plots. Most of the parameters show similar patterns as Figure 4, which indicate that those parameters are mixed well. But some SNP effects do not mix so well, which require larger number of MCMC loops and the bigger thinned number. In the following analysis, we still used those estimates, although they are not ideal to use.

#### Crohn’s Disease (CD) data set

First, we randomly selected one training data set out of 20 to compare the SNP effect estimation performance and the associated SNP detection performance of BayesRB, BayesR and logistic regression. Figure 18a shows that the BayesRB SNP effect estimates have a linear relationship with the BayesR estimates, except for two SNPs. The two SNPs are rs7593114 and rs6715049, both located on chromosome 2. Figure 18b compares the marginal logistic regression estimates to both BayesRB and BayesR estimates of the top 10 SNPs with the biggest BayesRB effects. Except for the two SNPs, BayesRB has slightly smaller estimates than marginal logistic regression, while BayesRB has much smaller estimates than the marginal logistic regression. The two SNPs which do not locate around the diagonal line are rs11887827 and rs6715049 on chromosome 2. Then, we generated LocusZoom plots [Pruim et al., 2010] to show the relationship of rs11887827, rs6715049 and rs7593114. Figure 19a shows that the three SNPs are highly correlated (*r*^2^ > 0.9), but while rs11887827 has a *p* value as small as 1.701 × 10^−5^, rs6715049 and rs7593114 do not have significant p values. Then, we conducted the logistic regression conditioning on rs11887827. Figure 19b shows that with rs11887827 conditioned on, rs6715049 and rs7593114 are pumped up with p values as small as 7.188 × 10^−20^ and 9.189 × 10^−11^, respectively. We recompared the SNP effect estimates of BayesRB to logistic regression, but with rs11887827 conditioned on. Figure 18c shows that all the top 10 SNPs’ BayesRB estimates are close to the logistic regression estimates, but slightly smaller than the logistic regression estimates. Then, we plotted manhattan plots based on the p values from both marginal logistic regression (Figure 20a, b) and logistic regression with rs11887827 conditioned on (Figure 20c, d). We explored the corresponding p values from logistic regression of the top 10 SNPs with the biggest BayesRB estimates (Figure 20a, c) and the biggest BayesR estimates (Figure 20b, d). Most of the top 10 SNPs locate on the top of the manhattan tower. Before rs11887827 is conditioned on, rs7593114 and rs6715049 locate at the bottom of the manhattan tower. After the rs11887827 is conditioned on, rs7593114 locates on the top of the manhattan tower, while rs6715049 is halfway up the tower. Table 2a shows that while rs11887827 and rs7593114 have perfect LD in the controls, 115 off diagonal entries in the cases. Table 2b shows that while only 5 out of 2225 controls having homozygous rs11887827 mutation and heterozygous rs6715049 mutation, 114 out of 1260 cases having this pattern. In Figure 20c, d, it seems that another SNP rs903228 is also halfway up the same tower of rs11887827. But actually, this SNP is 2,797kb away from rs11887827. We also investigated manhattan towers on chromosome 1 and 16. In Figure 20c, rs7515029 on chromosome 1 does not locate on the top of the manhattan tower. The SNP on the top of the tower is rs2201841, which is one of the top 30 biggest BayesRB estimated SNPs. rs7515029 and rs2201841 have low correlation (Figure 21a). In Figure 20c and d, two SNPs on the same tower in chromosome 16 have large BayesRB and BayesR estimates. But according to the Figure 21b, although the two SNPs are close to each other, they locate on different genes with a low correlation (*r*^2^ < 0.2). Therefore, it is reasonable that BayesRB and BayesR generate big estimates for both of the SNPs.

**Figure 18:**
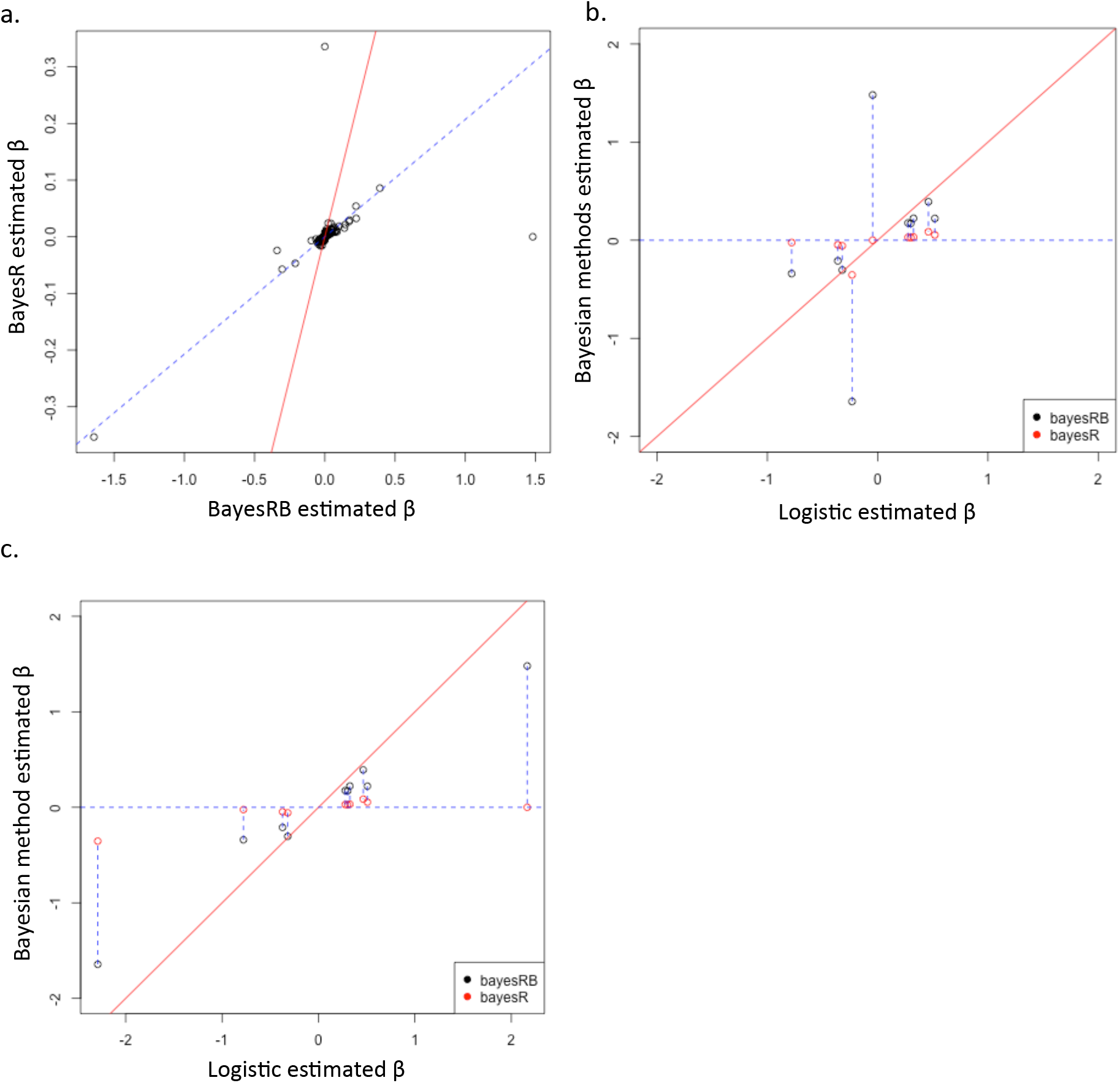
The comparisons of SNP effect estimates by BayesRB, BayesR, and logistic regression in one of the CD training data sets. The red solid lines are the diagonal lines. **a.** Compare the SNP effect estimates of BayesRB to BayesR. The blue dotted line is the fitted regression line, excluding the estimates of rs6715049 and rs7593114. **b.** Compare the marginal logistic regression estimates to BayesRB and BayesR estimates of the top 10 SNPs which have the biggest BayesRB estimates. **c.** Compare the logistic regression estimates conditioning on rs11887827 to BayesRB and BayesR estimates of the top 10 SNPs which have the biggest BayesRB estimates.

**Figure 19:**
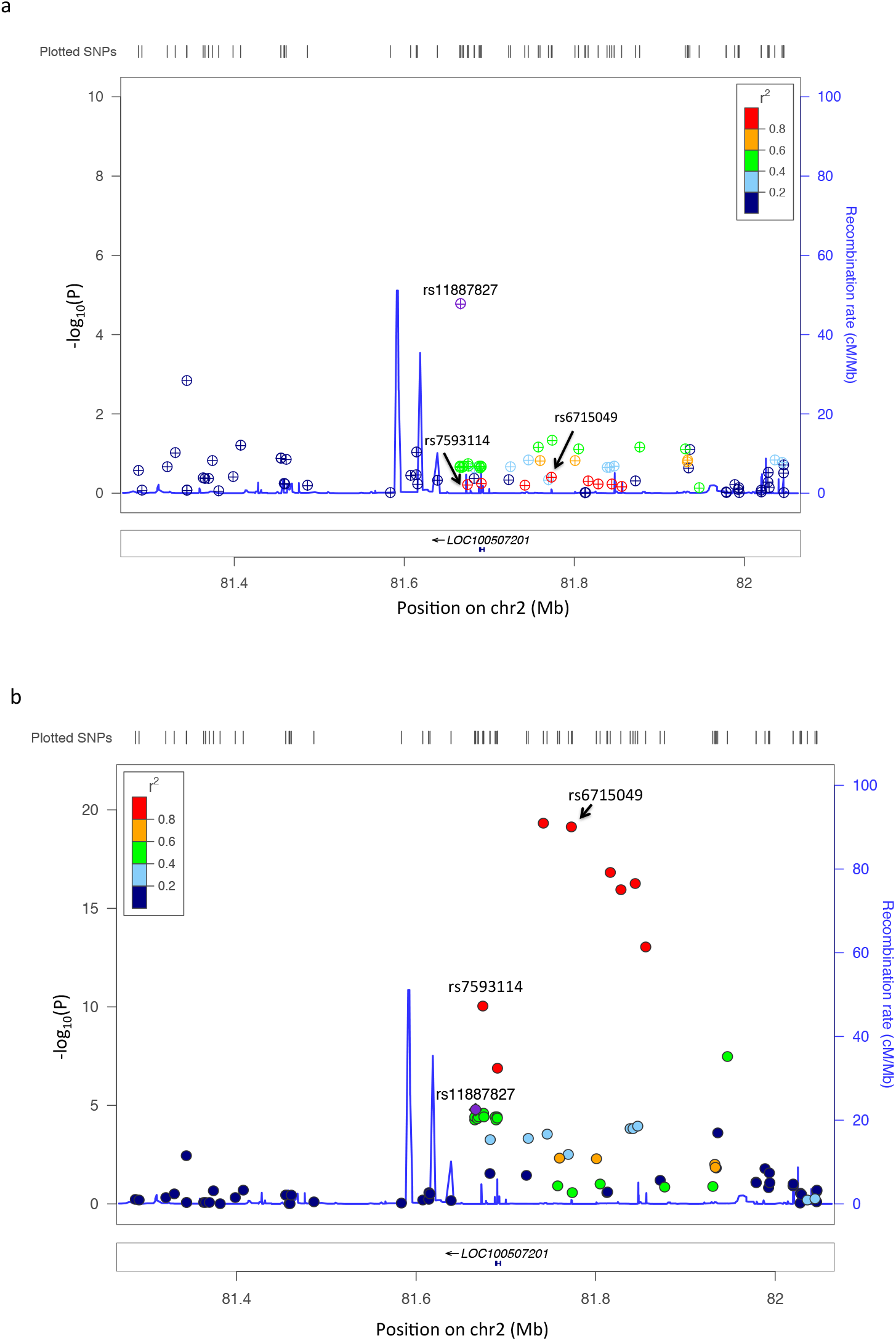
LocusZoom plots of 400kb region on chromosome 2, centered on rs11887827. **a.** The p values are generated from marginal logistic regression analysis. **b.** The p values are generated from logistic regression analysis with rs11887827 conditioned on.

**Figure 20:**
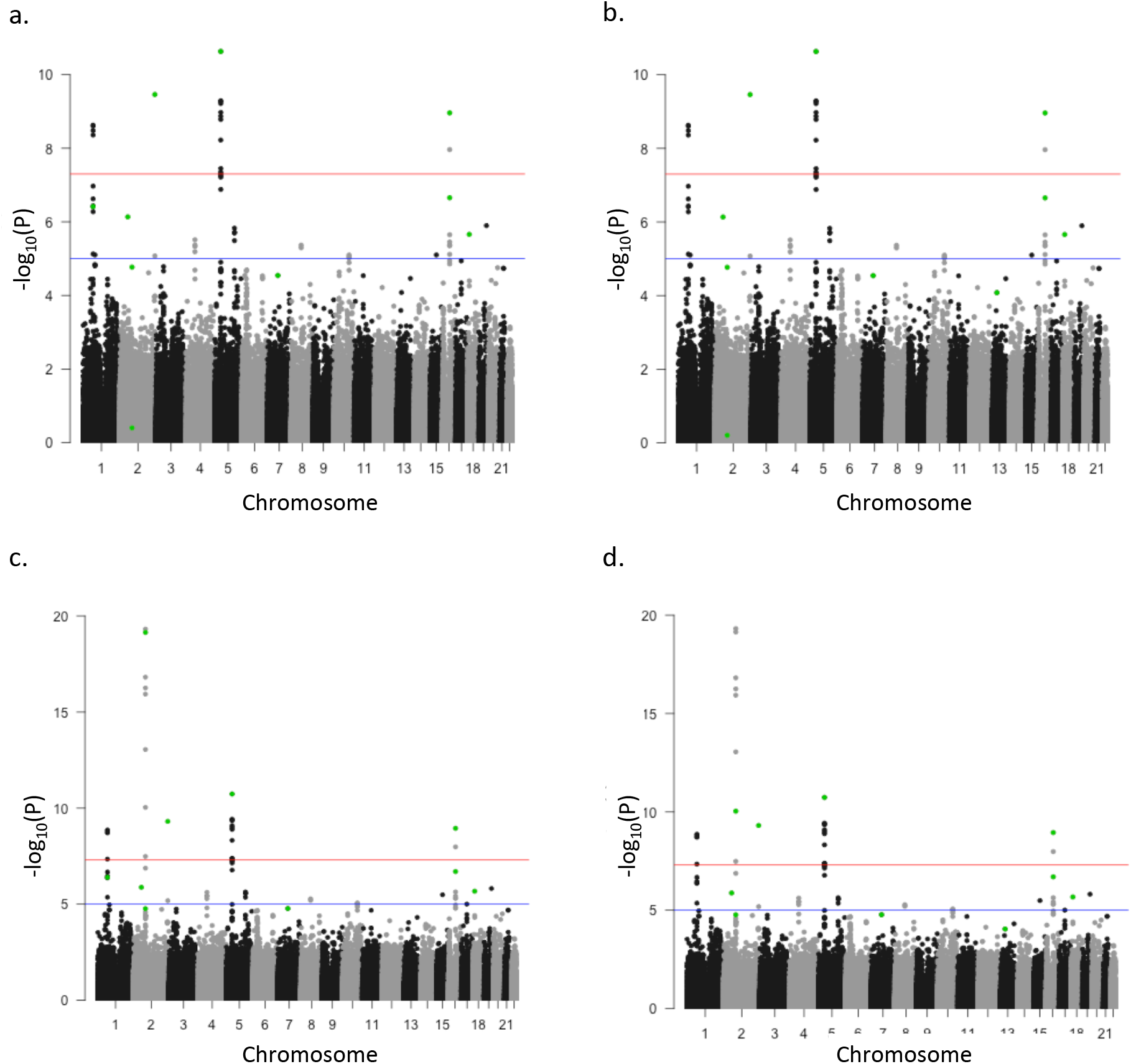
Manhattan plots of one of the CD training data sets before and after the SNP rs11887827 is conditional on with the top 10 BayesRB and BayesR estimated SNPs highlighted, respectively. **a.** The y axis is the −*log*_10_ of the marginal p values. The green dots indicate the 10 SNPs with the biggest BayesRB SNP effect estimates. **b.** Same manhattan plot as in **a**. The green dots indicate the 10 SNPs with the biggest BayesR SNP effect estimates. **c.** The y axis is the −*log*_10_ of the p values conditioned on rs11887827. The green dots indicate the 10 SNPs with the biggest BayesRB SNP effect estimates. **d.** Same manhattan plot as in **c**. The green dots indicate the 10 SNPs with the biggest BayesR SNP effect estimates.

**Table 2:**
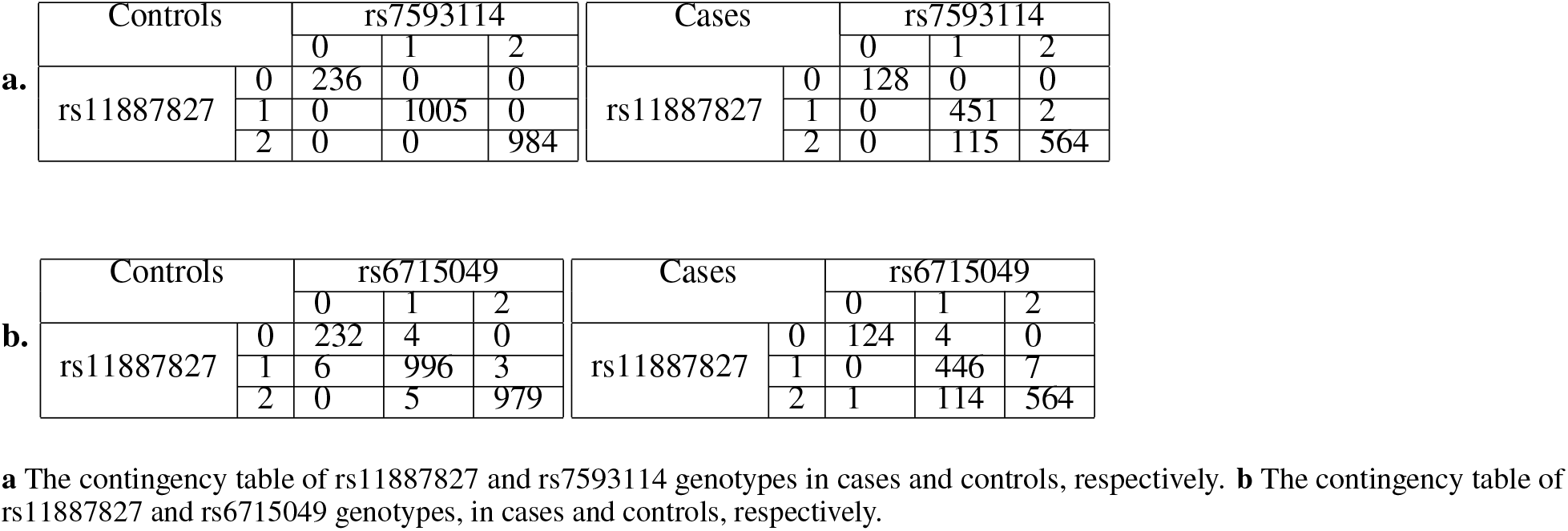
The contingency table of genotypes of rs11887827 and its highly correlated SNPs in cases and controls, respectively.

**Figure 21:**
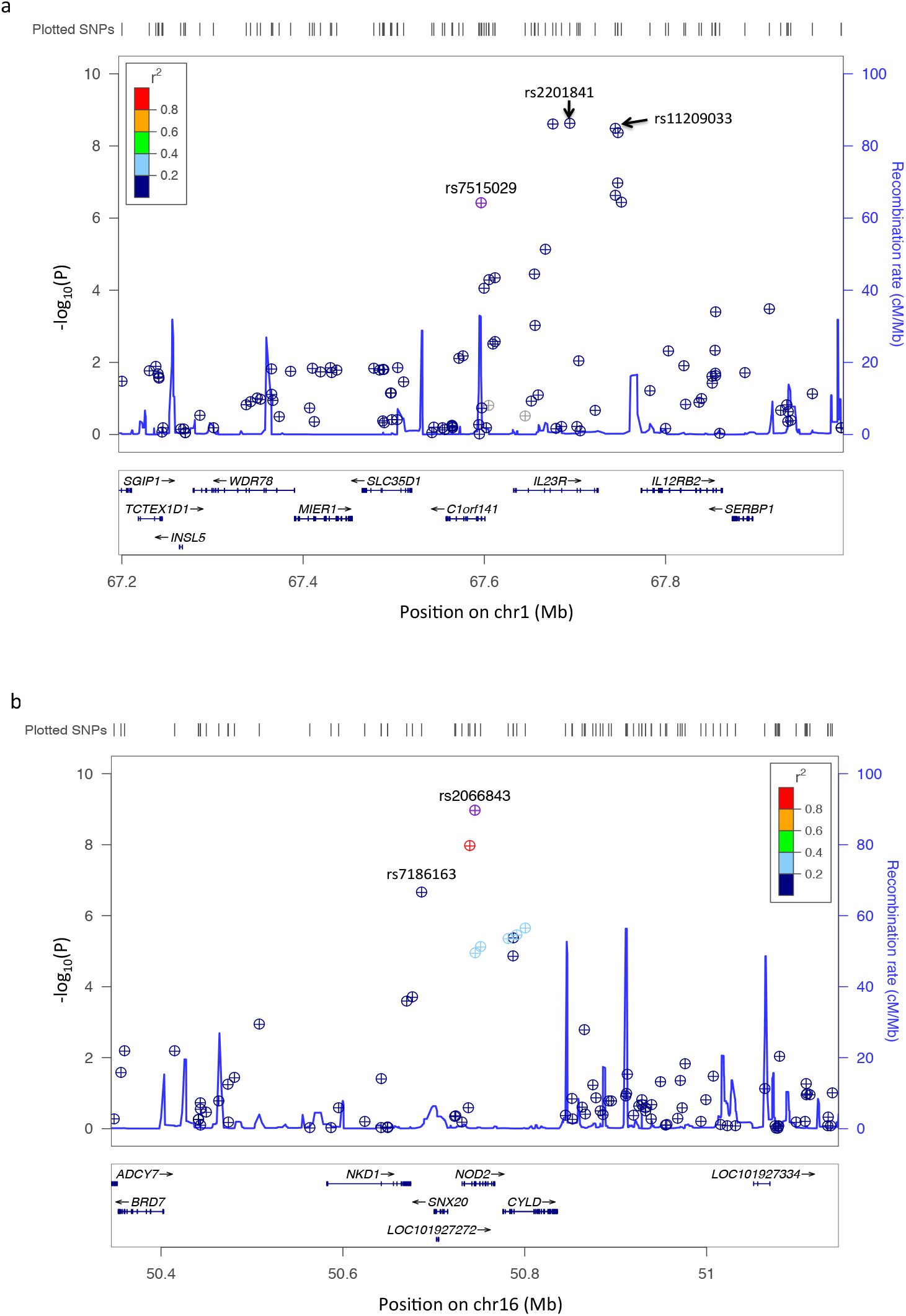
LocusZoom plots of 400kb region, centered on rs7515029 and rs2066843, respectively. The purple dot indicate the SNP that the plot is centered on. The p values are generated by marginal logistic regression. **a.** LocusZoom plot of 400kb region on chromosome 1, centered on rs7515029. **b.** LocusZoom plot of 400kb region on chromosome 16, centered on rs2066843.

We calculated the TPR and FPR to detect the 250kb windows containing the 201 previously reported SNPs. The 201 previously reported SNPs are obtained from GWAS Catalog [Welter et al., 2014] and Liu at al. [2015]. Only the SNPs reported by studies with European ancestry samples are included. All 201 SNPs have reported p values smaller than 5 × 10^−8^. Figure 22a and b show that logistic regression has the biggest TPRs, followed by LASSO, while BayesR has the lowest TPR, under the same value of FPRs. When the FPR is 0.08, the TPR is around 0.2 for all the methods.

**Figure 22:**
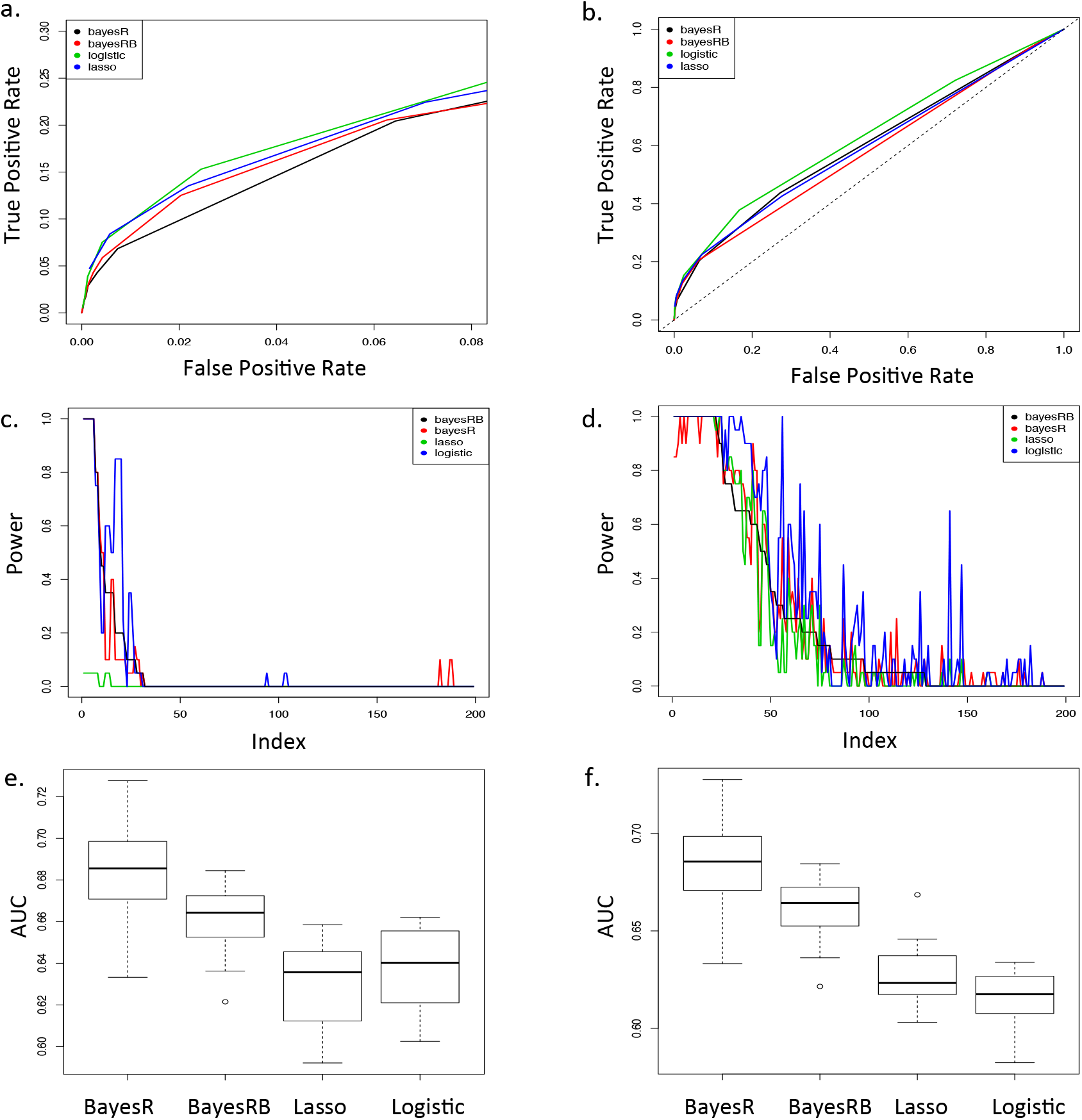
Comparisons of BayesRB, BayesR, logistic regression and LASSO’s associated SNP selection performance and risk prediction performance in the CD data sets. **a.** and **b.** True positive rate vs. false positive rate of detecting the 250kb windows containing the 201 previously reported SNPs. The black dotted line in **b.** is the diagonal line. **c.** and **d.** Power to detect the windows containing the 201 previously reported CD associated SNPs in the 20 replicates. The SNPs are sorted by a decreasing order of their BayesRB powers. **e.** and **f.** AUCs of BayesRB, BayesR, LASSO, and logistic regression in 20 replicates. In **c** and **e**, The thresholds of logistic regression and the LASSO are set under the FPR of 0.001. In **d** and **f**, The thresholds of logistic regression and the LASSO are set under the FPR of 0.05.

Then, we compared the power of BayesR, BayesRB, logistic regression and LASSO to have each window containing causal SNPs identified within the 20 replicates. Figure 22c and d show that using the thresholds under the same FPR of both 0.001 and 0.05, logistic regression has highest power to detect the previously reported SNPs, and LASSO has the lowest power. For some SNPs, BayesRB has bigger power than BayesR, but for other SNPs, it does not.

At last, we compared the performance of risk prediction of BayesR, BayesRB, logistic regression and LASSO on the testing data set by comparing the AUCs. Figure 22e and f show that BayesRB and BayesR generate bigger AUCs than LASSO and logistic regression. But BayesR’s median value of AUC is slightly higher than BayesRB. BayesRB’s prediction is more precise than BayesR.

#### Bipolar Disorder (BD) data set

For Bipolar Disorder (BD) data set, we also randomly selected one training data set out of 20 to compare the SNP effect estimation performance and the associated SNP detection performance of BayesRB, BayesR and logistic regression. Figure 23a shows that the BayesRB SNP effect estimates have a linear relationship of the BayesR estimates, except for two SNPs: rs4923955 and rs16957168. And the other two SNPs have much bigger BayesRB and BayesR SNP effect estimates than other SNPs. The two SNPs are rs12050604 and rs1381855. The four SNPs locate close to each others on chromosome 15. The farthest two SNPs are only 62.3kb away from each other. Among the four SNPs, only rs12050604 has a significant *p* value (5.08 × 10^−7^) in marginal logistic regression analysis. The other three SNPs all have *p* value bigger than 0.05. The two green dots locating at the bottom in chromosome 15 in Figure 24a are rs4923955 and rs1381855, which do not have significant *p* value but has top 10 SNP effect estimates of BayesRB. The two green dots locating at the bottom in chromosome 15 in Figure 24b are rs16957168 and rs1381855, which do not have significant p values but have top 10 SNP effect estimates of BayesR. However, when we reestimated the SNP effects using logistic regression conditional on rs12050604, both rs1381855 and rs16957168 are pumped up (Figure 25). Figure 24c and d show that with the rs12050604 conditioned on, while both rs12050604 and rs1381855 are on the top of the manhattan tower, rs4923955 is still at the bottom, and rs16957168 is halfway up the tower. Figure 23b and c show that before the rs12050604 is conditioned on, rs12050604, rs1381855 and rs4923955 have much larger BayesRB effect estimates than logistic regression; however, after rs12050604 is conditioned on, BayesRB has similar SNP effect estimates of rs12050604 and rs1381855 to logistic regression estimates, but it still has much larger effect estimate of rs4923955 than logistic regression. Table 3 shows the contingency table of rs12050604 and the other three SNPs’ genotypes. While only 14 out of 2224 controls have homozygous rs16957168 mutations and heterozygous rs12050604 mutations, 374 out of 1332 cases have the same genotypes. While 193 out of 2224 controls have homozygous rs12050604 mutations and heterozygous rs4923955 mutations, only 19 out of 1332 cases have the same genotypes. The joint distributions of rs12050604 and rs1381855 genotypes are similar in cases and controls.

**Figure 23:**
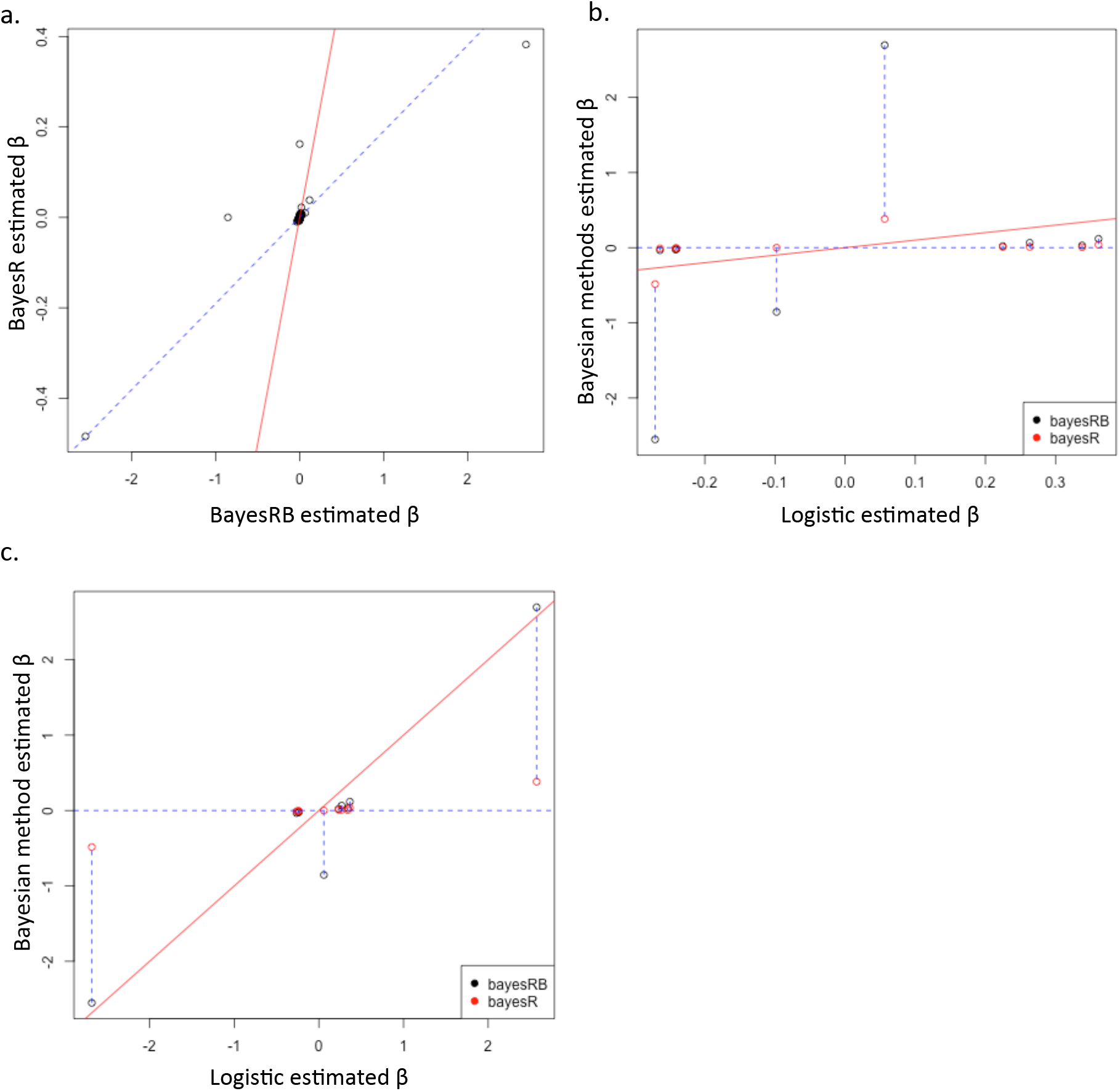
The comparisons of SNP effect estimates by BayesRB, BayesR, and logistic regression in the BD data sets. The red solid lines are the diagonal lines. **a.** Compare the SNP effect estimates of BayesRB to BayesR. The blue dotted line is the fitted regression line, excluding the estimates of rs1381855 and rs4923955. **b.** Compare the marginal logistic regression estimates to BayesRB and BayesR estimates of the top 10 SNPs which have the biggest BayesRB estimates. **c.** Compare the logistic regression estimates conditional on rs12050604 to BayesRB and BayesR estimates of the top 10 SNPs which have the biggest BayesRB estimates.

**Figure 24:**
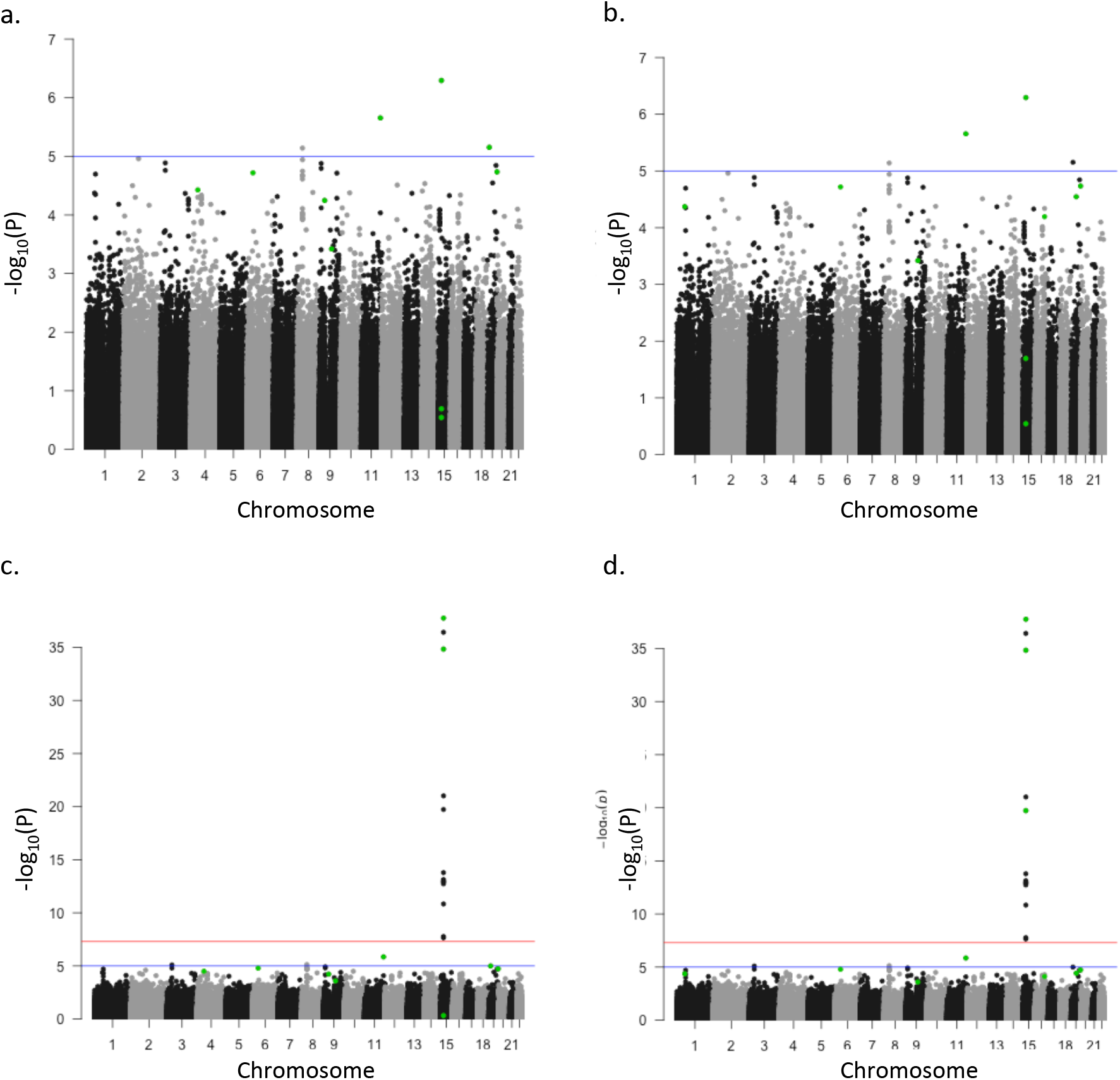
Manhattan plots of one of the BD training data sets before and after the SNP rs12050604 is conditioned on with the top 10 BayesRB and BayesR estimated SNPs highlighted, respectively. **a.** The y axis is the *-log*_10_ of the marginal p values of all the SNPs. The green dots indicate the 10 SNPs with the biggest BayesRB SNP effect estimates. **b.** Same manhattan plot in **a**. The green dots indicate the 10 SNPs with the biggest BayesR SNP effect estimates. **c.** The y axis is the −*log*_10_ of the p values of all the SNPs conditioned on rs12050604. The green dots indicate the 10 SNPs with the biggest BayesRB SNP effect estimates. **d.** Same manhattan plot in **c**. The green dots indicate the 10 SNPs with the biggest BayesR SNP effect estimates.

**Figure 25:**
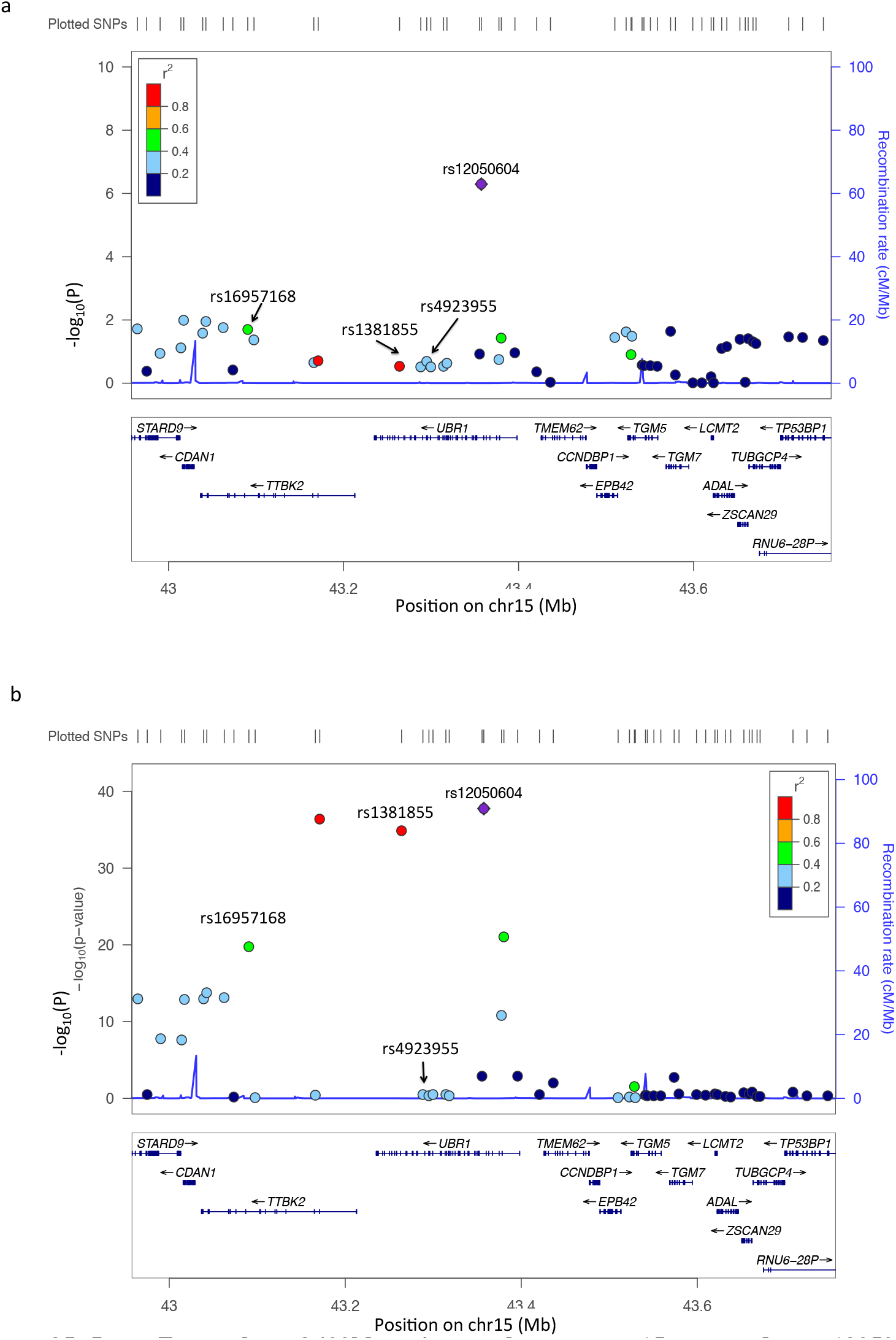
LocusZoom plots of 400kb region on chromosome 15, centered on rs12050604. **a.** The p values are generated from marginal logistic regression analysis. **b.** The p values are generated from logistic regression analysis with rs12050604 conditioned on.

**Table 3:**
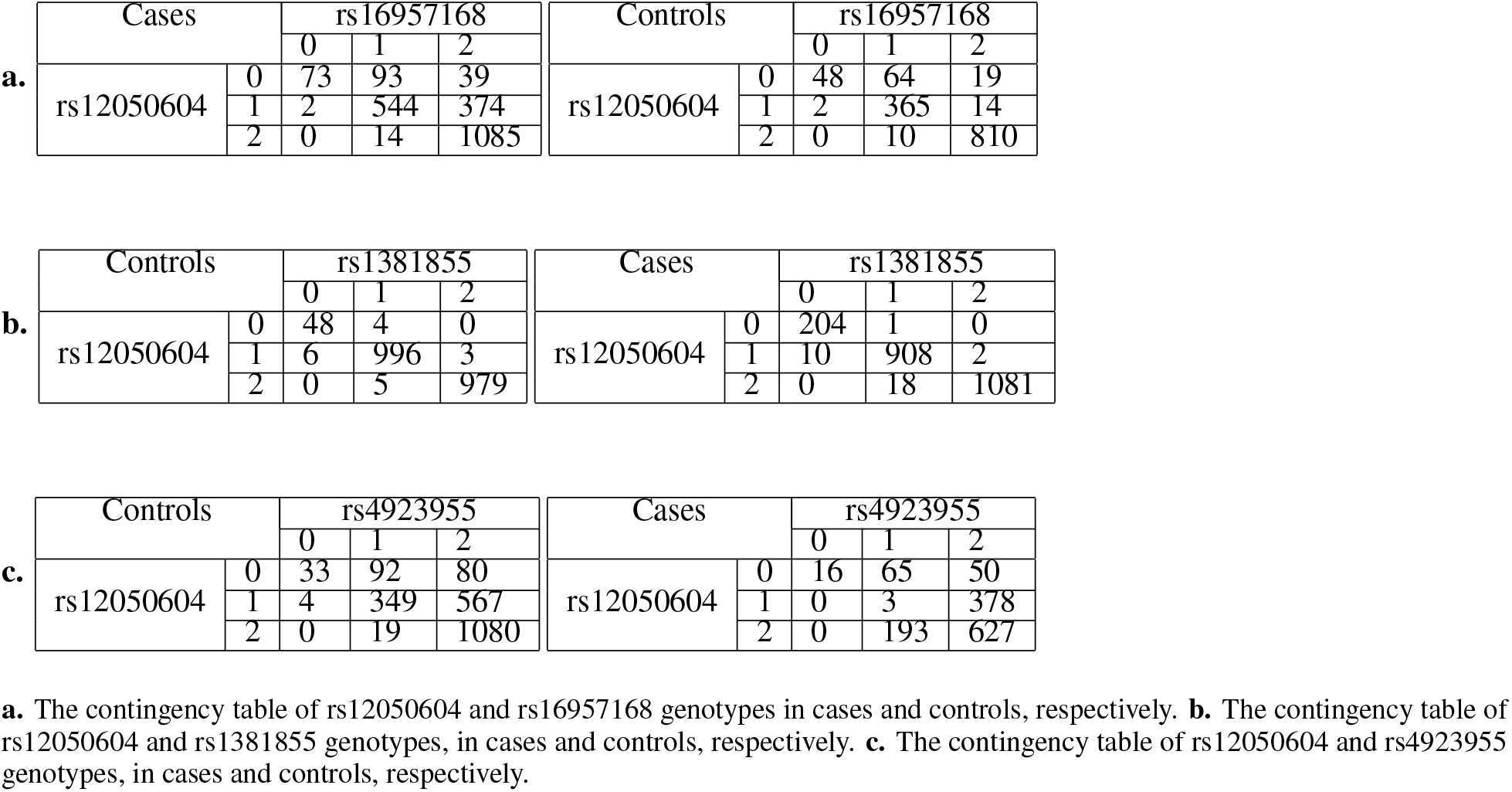
The contingency table of genotypes of rs12050604 and its highly correlated SNPs in cases and controls, respectively.

For the BD data set, we also compared the TPR and FPR of BayesRB, BayesR, logistic regression, and LASSO to detect the 250kb windows containing the previously reported SNPs; compared the power of the four methods of having each window containing causal SNPs identified within the 20 replicates; and compared the performance of risk prediction of the four methods on the testing data set. The 497 previously reported SNPs are obtained from GWAS catalog [Welter et al., 2014] with reported p values smaller than 5 × 10^−8^. Only the SNPs reported by studies with European ancestry samples are included. Figure 22a shows that when the FPR is smaller than 0.08, under the same value of FPRs, logistic regression and LASSO have similar TPRs; BayesRB and BayesR have similar TPRs. The logistic regression and LASSO have higher TPR than BayesRB and BayesR under the same value of FPRs. When the FPR is 0.08, the TPR is only around 0.1 for all the methods. 22b shows that, all in all, the TPRs are similar for the four methods, but BayesRB has slightly lower TPR than the other three under the same FPR. Figure 22c and d illustrate that using the thresholds under the FPR of both 0.001 and 0.05, the four methods have similar powers. Figure 22e and f show that BayesRB and BayesR generate bigger AUCs than LASSO and logistic regression. But BayesR’s median value of AUCs is higher than BayesRB. BayesRB and BayesR have similar precise predictions.

## 4 Discussion

We developed a Bayesian method called BayesRB for dichotomized phenotypes to select associated SNPs, estimate the SNP effects, and make the prediction of the disease risks, taking account of the effects of SNPs in the whole genome. We also evaluated BayesRB performance and compared the performance with BayesR, LASSO, and logistic regression by using pilot simulated data sets, 50 genome-wide simulated data sets, and WTCCC data sets. The results showed that BayesRB was a promising method for dichotomized disease/trait prediction.

When developing our BayesRB method, we investigated two implementations. In the first, we estimated the genetic variance 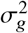, similar to what Moser et al. [2015] did. We set a uniform prior for the 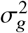 and estimated it using Metropolis-Hasting sampling. In the second, we fixed the 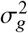 as Erbe et al. [2012] did. Based on our results comparing these two approaches, we decided not to use Metropolis-Hasting sampling to estimate 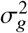 for the realistic pilot simulated data set, genome-wide simulated data sets, and the real data sets for the following two reasons: First, a fixed 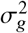 generates similar ***β*** estimates as the random 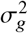 does. The unrealistic data set showed that when 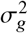 is set to a fixed value bigger than 10 or is estimated using Metropolis-Hasting sampling, the BayesRB ***β*** estimates are almost equivalent to each other (Figure 6). Using the Metropolis-Hasting sampling, the variance of the proposal distribution has to be tuned manually, which has low efficiency for large data sets. So, it is reasonable to use fixed 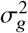 instead. Second, the variance of the SNP effects of the realistic data set, genome-wide simulated data sets, and the real data sets are all very small compared to the individual variances *λ*. The four categories are distinguished by the four different variances of the normal distributions. Only when the variance is big enough, the four categories can be distinguished from each other (Figure 27a). When the variance of the SNP effects are very small compared to the *λ*, it is hard to tell the difference of the four categories (Figure 27b). In this case, 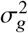 cannot be estimated well. Therefore, we set 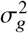 fixed for the realistic data set of the pilot simulated data sets, genome-wide simulated data sets, and the real data sets.

**Figure 26:**
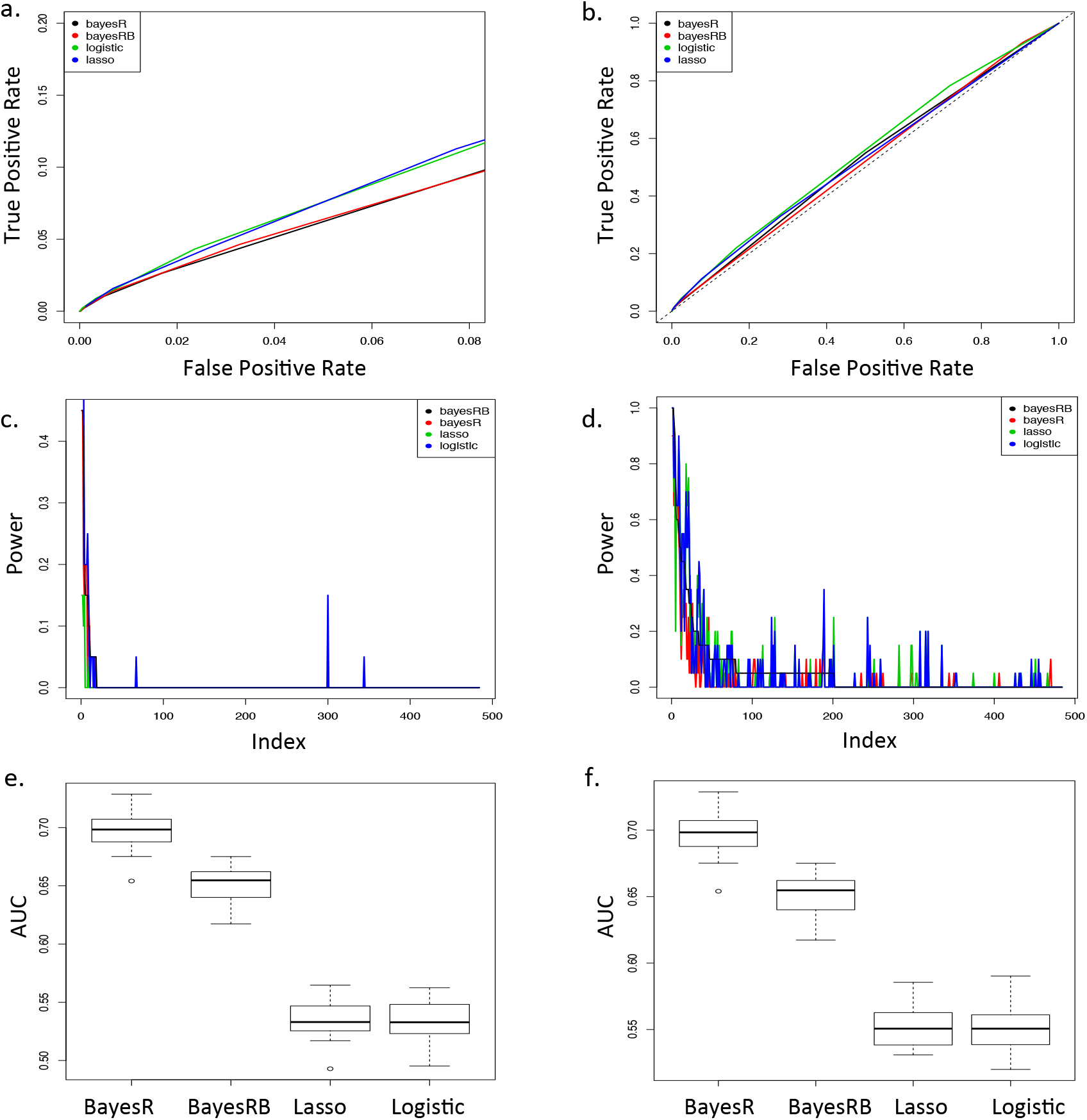
Detection of BayesRB’s associated SNP selection performance and risk prediction performance in the BD data sets. **a.** and **b.** True positive rate vs. false positive rate of detecting the 250kb windows containing the 497 previously reported SNPs. The black dotted line in **b.** is the diagonal line. **c.** and **d.** Power to detect the windows containing the 497 previously reported BD associated SNPs in the 20 replicates. The SNPs are sorted by a decreasing order of their BayesRB powers. **e.** and **f.** AUCs of BayesRB, BayesR, LASSO, and logistic regression in 20 replicates. In **c** and **e**, The thresholds of logistic regression and the LASSO are set under the FPR of 0.001. In **d** and **f**, The thresholds of logistic regression and the LASSO are set under the FPR of 0.05.

**Figure 27:**
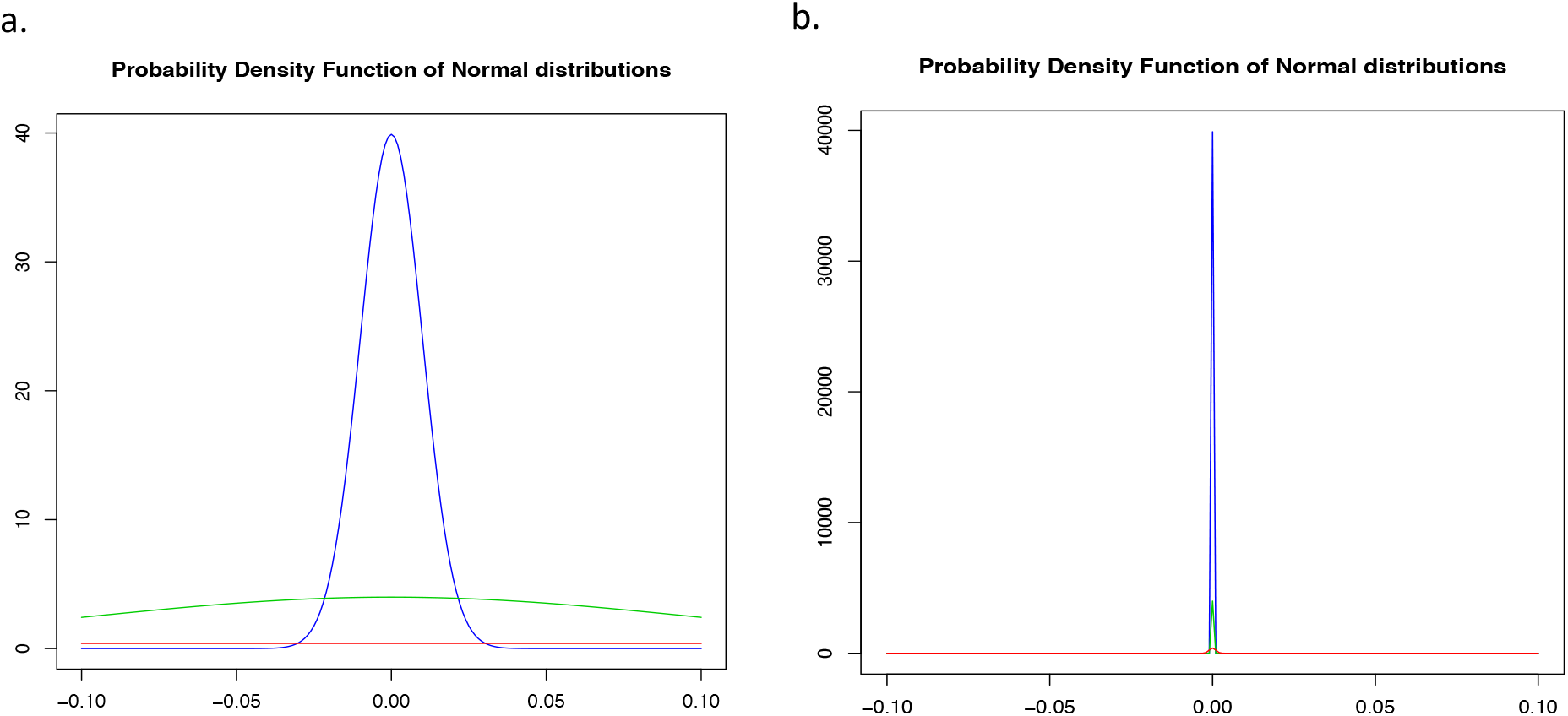
The probability density functions of the normal distributions. The blue curves represent the 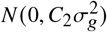; the green curves represent the 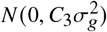; and the red curves represent the 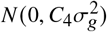, where (*C*_2_, *C*_3_, *C*_4_) = (10^−4^, 10^−3^, 10^−2^), respectively. **a.** 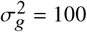. **b.** 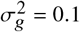.

When the 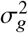 is small 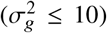, the ***β*** is underestimated. Both the mean and the variance of the posterior distribution of ***β*** is a function of 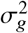. When the 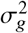 is small, even the SNP is assigned to the fourth category, the ***β*** prior has a small variance and a zero mean and the ***β*** prior has a large effect on the ***β*** posterior distribution according to the Formula 3.14. So the BayesRB ***β*** tends to be underestimated. When the 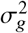 is large enough 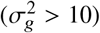, the ***β*** prior does not have a large effect on the ***β*** posterior distribution any more. Data then drives the ***β*** posterior distribution.

When the 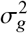 is large enough 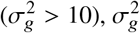 affects the SNP categorization, thus, affects the shrinkage level of the SNP effect estimation. When the 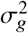 is larger, the SNPs are more likely to be assigned to the categories with smaller variance and a larger proportion of SNPs are assigned to the category 1, so the BayesRB shrinks the SNP effect more.

We examined the SNP effect estimation performance of BayesRB, BayesR, and logistic regression. LASSO underestimates the SNP effects [Wu et al., 2009], therefore, it is not included in the comparison. The results in both genome-wide simulated data sets and the real data sets show that for the large effect SNPs, BayesRB provides similar SNP effect estimates to the logistic regression estimates, which is unbiased; while for the small effect SNPs, BayesRB shrinks their SNP effects to zero (Figure 11, Figure 18, Figure 23). BayesR underestimated the SNP effects. This result is consistent with Gianola et al. [2013].

In the CD data analysis, from the BayesRB and BayesR SNP effect estimation results, we discovered that the joint distributions of rs11887827 and two of its highly correlated SNPs are quite different in cases and in controls (Table 2); in the BD data analysis, we discovered that the joint distributions of rs12050604 and two of its highly correlated SNPs are quite different in cases and in controls (Table 3). In the WTCCC study, the case data sets and the control data sets were generated separately, according to the WTCCC website. We made scatter plots using the fluorescent signal intensity data of the two alleles of rs11887827 and rs12050604 for each individual, as well as their highly correlated SNPs. Intensity plots of both rs11887827 and rs12050604 show four clusters instead of three (Figure 28), which may be caused by duplications [Kumasaka et al., 2011]. All the highly correlated SNPs show a normal pattern of three clusters (data not shown). Figure 28 show that, for both rs11887827 and rs12050604, different decisions were made in controls and cases when calling the genotypes in cluster 2. For both SNPs, controls in cluster 2 are treated as having 1/0 genotypes, but cases in cluster 2 are treated as having 1/1 genotypes. Thus, different decisions on how to call the genotypes in cluster 2 lead to the different joint distributions of genotypes in cases and controls.

**Figure 28:**
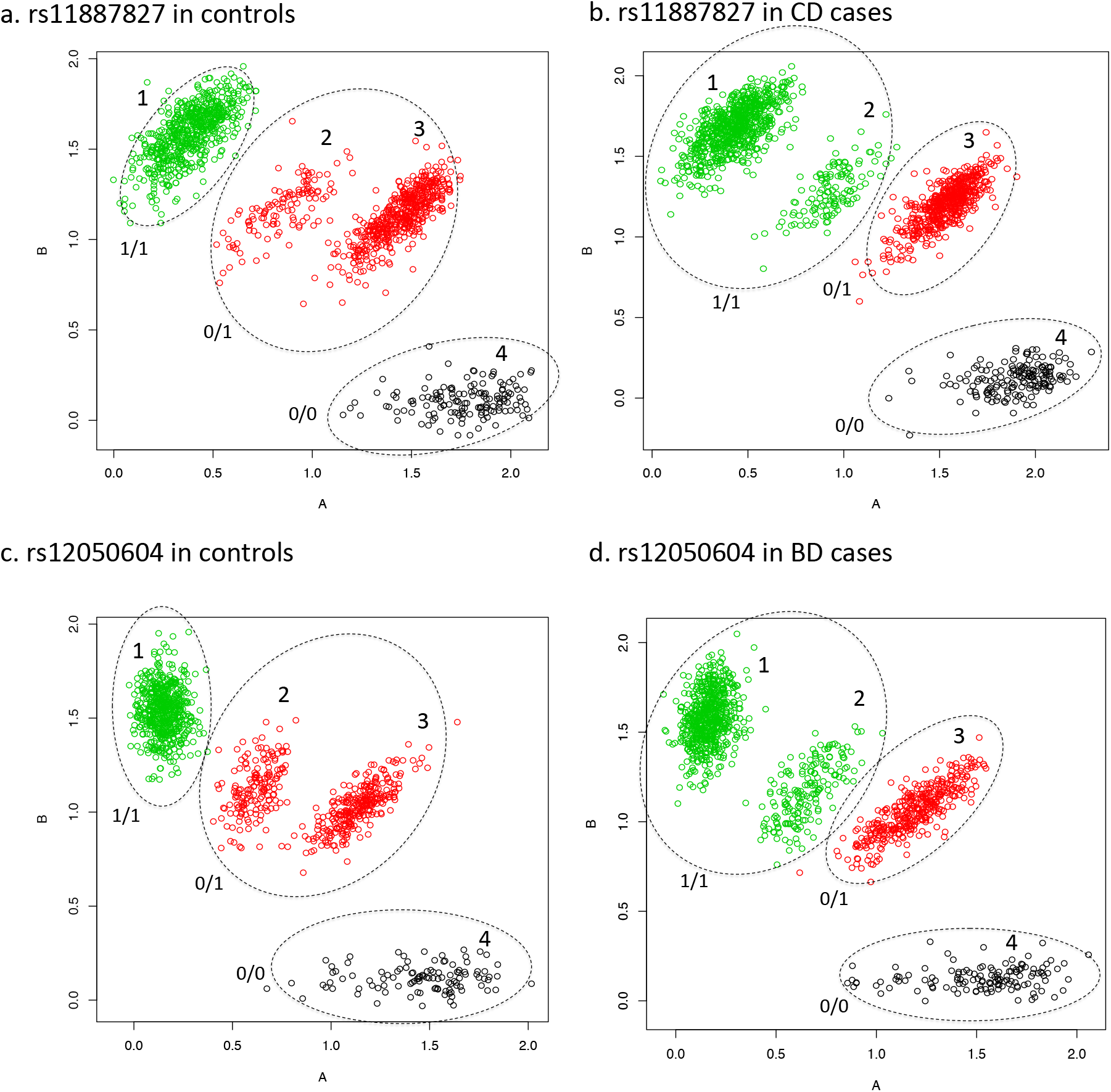
Fluorescent signal intensity plots for rs11887827 and rs12050604 in controls and cases, respectively. The x-axis corresponds to the allele A and the y-axis corresponds to the allele B, respectively. The black dots indicate the individuals with coded genotypes of 0/0. The red dots indicate the individuals with coded genotypes of 0/1. The green dots indicate the individuals with coded genotypes of 1/1. The numbers “1”, “2”, “3” and “4” in the plots indicate the four clusters.

The observation discussed above in the CD and BD data sets are due to batch effects, however, similar observation could be due to some real signals. In reality, it is possible that some markers become significant after conditioning on other markers. It is also possible that some markers are no longer significant after conditioning on other markers. In GWAS, people often conduct an additional conditional analysis to further investigate the markers conditional on the significant ones. If the first marker itself is not significant (like rs11887827 has *p* value < 10^−5^ in CD) in marginal logistic regression, then this marker will not be conditioned on, and thus no other markers can be discovered. BayesRB estimates the SNP effects simultaneously and makes up the defects of the conventional marginal logistic regression method. It can save effort to conduct the conditional analysis after discovering the significant markers. It may not fail to detect the jointly significant markers, nor mistakenly include the markers in the model that are significant only because they are in LD with more significant markers. In addition, BayesRB can help with quality control. For example, in our analyses, it discovered two SNPs affected by batch effects. These problematic SNPs should be deleted from the data sets. All in all, the above reasons illustrates the potential value of the BayesRB approach over the conventional marginal logistic regression method.

We examined the associated SNP selection abilities of BayesRB, BayesR, LASSO, and logistic regression, using both genome-wide simulated data sets (Figure 13) and the real data sets (Figure 22a,b and Figure 26a,b). In the genome-wide simulated data sets, we set the causal SNPs at the beginning of the simulation, therefore, the TPR to detect big effect causal SNPs can be as large as 1; and the TPR to detect medium effect causal SNPs can also be larger than 0.6, when the FPR is 0.08. But in the real data sets, it is unknown which SNPs are the true causal SNPs. So we calculated the TPR and FPR of detecting the previously reported SNPs. The TPR to detect the previously reported CD and BD SNPs are as low as around 0.2 and 0.1 respectively, when the FPR is 0.08. In addition, in the genome-wide simulated study, the proportion of windows containing causal SNPs being discovered in at least one replicates is much bigger than that in the real data study. The low TPRs of detecting the previously reported SNPs within 250kb regions, and the low power of having the windows containing the previously reported SNPs being detected in CD and BD data sets are because of the following reasons: First, for the real data sets, we calculated TPR and FPR using all of the previously reported SNPs regardless of how big their effect size estimates turned out to be in the BayesRB analysis. So, it is reasonable that the TRP is smaller than only using the big and medium effect SNPs as we did in genome-wide simulated data sets. Second, some previously reported associated SNPs are discovered from studies with large sample sizes (> 20,000). But the sample sizes in the CD and BD training data sets are only 3,485 and 3,556, respectively. Given the relatively small sample sizes in CD and BD data sets, only SNPs having relatively big odds ratios are detectable. Third, even if some SNPs that are discovered to be associated with CD and BD from data sets with similar sample sizes to those of the WTCCC CD and BD data sets, they may not be detectable using the WTCCC CD and BD data sets. Fourth, some SNPs are reported to be associated with CD or BD, but they may not be. The SNPs may be discovered due to some false positive signals, and thus cannot be replicated using other data sets.

Based on the results of TPR, FPR and power of identifying windows containing causal SNPs in the genome-wide simulated data sets (Figure 13), and identifying windows containing previously reported SNPs in the real data sets (Figure 22, Figure 26), we concluded that BayesRB does not perform better than the other three methods. However, BayesRB is still a promising method for identifying associated SNPs in real studies and guiding the direction of future studies. The BayesRB associated SNPs are those with proportion of iterations that are assigned to the category 2, 3 or 4 (assignment proportion) bigger than the threshold. Figure 7 and Figure 12 shows that the SNPs with bigger logistic regression estimates have bigger assignment proportions, and thus are likely to be selected as the BayesRB associated SNPs. This shows BayesRB’s good performance in terms of its ability to select associated SNPs. Figure 7 and Figure 12 are slightly different: First, many more SNPs in Figure 7 have big assignment proportions than that in Figure 12. This is because Figure 7 is based on the unrealistic data sets, which contains SNPs with bigger effects than the reality; while Figure 12 is based on the genome-wide simulated data sets, which contains the SNPs with effects similar to the real data sets. Second, in Figure 12, some SNPs have logistic estimates as big as around 0.4, but have assignment proportions close to 0; while in Figure 7, all the big logistic regression estimated SNPs have big assignment proportions. This is due to the reason that in the unrealistic data sets, all the SNPs are independent of each other; while in the genome-wide simulated data sets, the SNPs are not fully independent. The SNPs having big logistic regression estimates but small proportions in the genome-wide simulated data sets may be correlated to some other more significant SNPs, thus, they are not selected in the model. Table 4 shows the detailed information of the top 10 SNPs with the biggest assignment proportions in the CD data set, excluding the two SNPs on chromosome 2 affected by the batch effects. 8 out of 10 SNPs locate within the 500 kb windows of the previously reported CD associated SNPs. Table 5 shows the detailed information of the top 10 SNPs with the biggest assignment proportions in the BD data set, excluding the three SNPs on chromosome 15 affected by the batch effects. 7 out of 10 SNPs locate within the 500kb windows of the previously reported BD associated SNPs. While BayesRB can be applied to identify the associated SNPs in the real studies, it also suggests that, in the future, more studies need to be conducted to assess the two SNPs that were not previously reported to be associated with CD and the three SNPs that were not previously reported to be associated with BD.

**Table 4:**
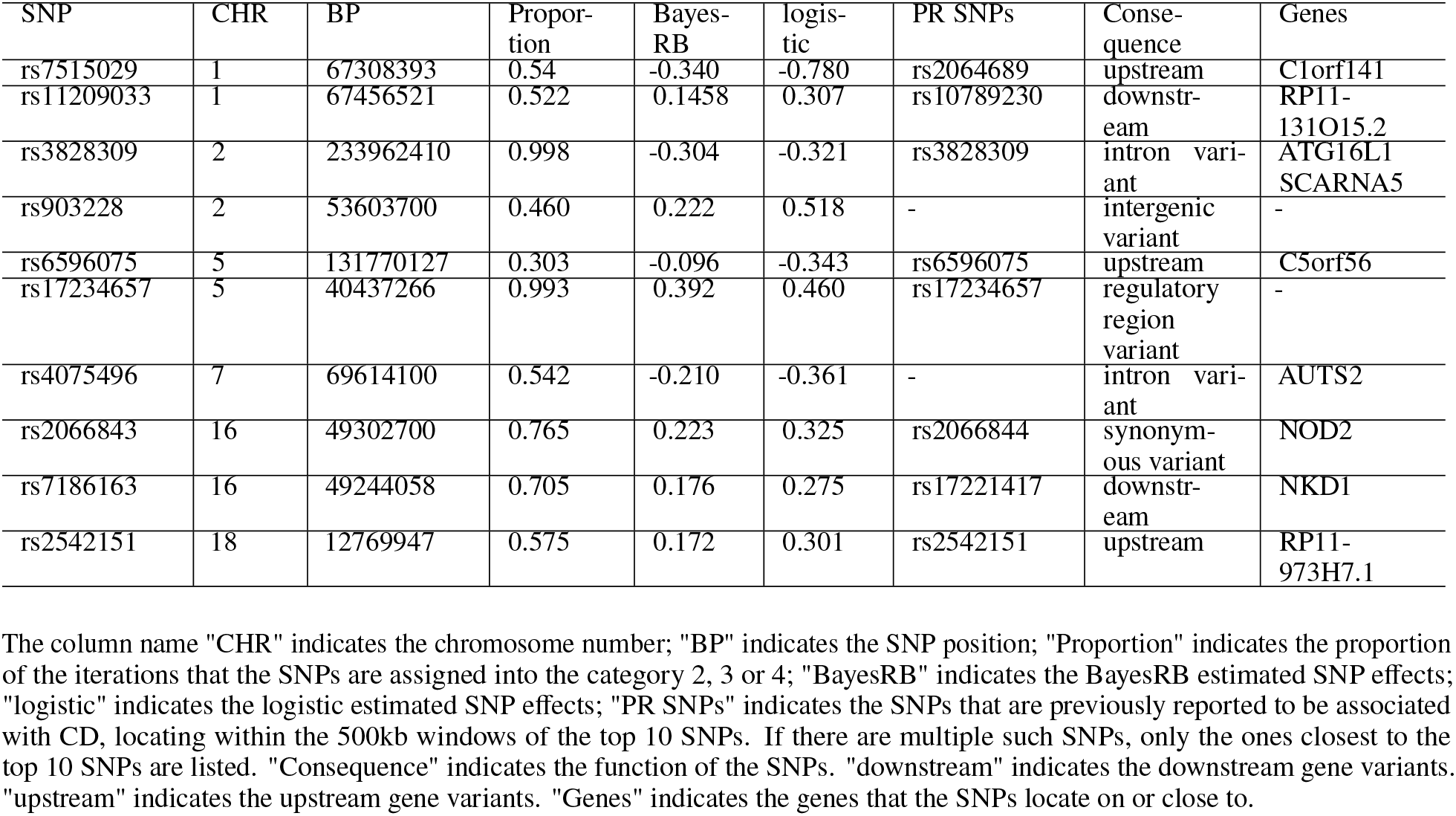
Detailed information of the top 10 SNPs with the biggest proportions to be assigned to the category 2, 3 or 4, excluding the two SNPs on chromosome 2 affected by the batch effects.

**Table 5:**
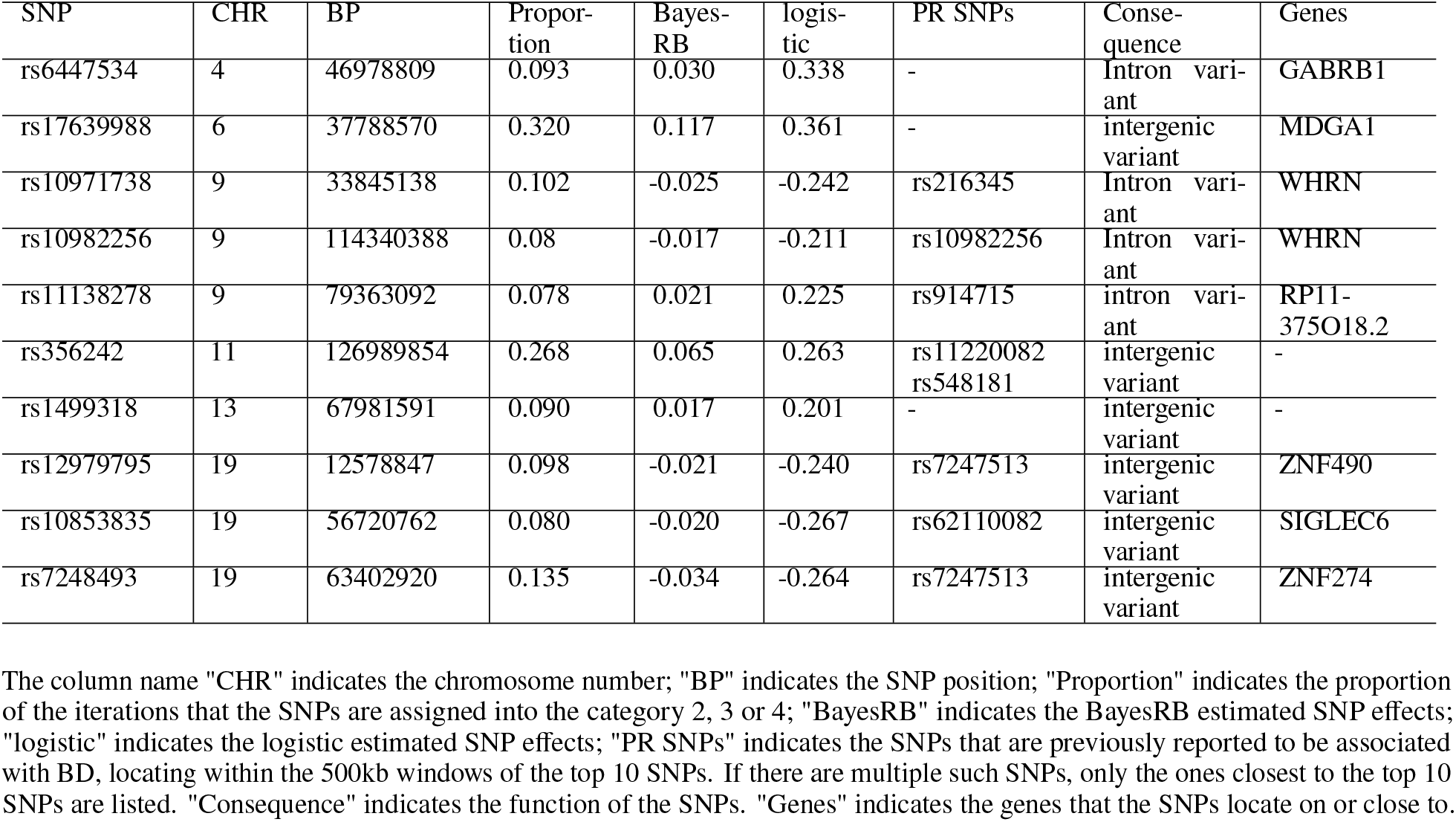
Detailed information of the top 10 SNPs with the biggest proportions to be assigned to the category 2, 3 or 4, excluding the three SNPs on chromosome 15 affected by the batch effects.

We also measured the prediction performance of BayesRB, BayesR, LASSO, and logistic regression using AUCs. BayesR has the best prediction performance than the other three methods, followed by BayesRB. In the genome-wide simulated data sets, which are not affected by the batch effects, when the FPRs of logistic regression and LASSO are 0.001, BayesR and BayesRB have slightly bigger AUCs than logistic regression and LASSO (Figure 16). In the CD and BD data sets, BayesR and BayesRB have much larger AUCs than logistic regression and LASSO (Figure 22e,f and Figure 26e,f). One possible reason is that in CD and BD data sets, the BayesRB and BayesR models not only contain rs11887827 and rs12050604, respectively, but also contain one of their correlated SNPs. Containing rs11887827 and rs12050604, respectively, in the models alone, as logistic regression does, does not fully account for the association signal in CD and BD. Therefore, BayesRB and BayesR have much better risk prediction performance than logistic regression and LASSO. The AUCs should be re-compared after more thorough quality control to identify and delete the SNPs affected by batch effects. But if the data truly contains some jointly significant markers, BayesRB and BayesR are expected to show better prediction performance than the conventional approaches.

When calculating the power of identifying the windows that containing the causal SNPs and calculating the AUCs of logistic regression and LASSO, we use the thresholds with the FPR equals to 0.001. A multiple comparison adjusted FPR is more appropriate (~ 10^−6^), but we did not use the multiple comparison adjusted FPR in this study. The reason is justified below. When the FPR is constrained to 0.001, the power of identifying most of the windows containing the causal SNPs are small. If the FPR were smaller, less SNPs would be identified as associated SNPs. Thus, the power of identifying the windows that containing the causal SNPs will be even smaller. In this case, all the four methods performs equally poor. When FPR is constrained to 0.001, four methods’ associated SNP selection performance can be better compared with each other. Considering the consistency of the dissertation, when calculating the AUCs of logistic regression and LASSO, we also set the thresholds using a FPR of 0.001.

It is recommended to use BayesRB instead of BayesR for dichotomous outcome data, even though BayesR has slightly better risk prediction performance than BayesRB. First, the SNP odds ratios can be estimated by BayesRB, but cannot be estimated by BayesR. BayesRB is based on a logistic regression model, while BayesR is based on an ordinary linear model. So, the estimated SNP odds ratios can be calculated by taking the log of the estimated SNP effects of BayesRB, but cannot be calculated by the estimated SNP effect of BayesR. Second, BayesRB can predict the risk of getting disease, but BayesR cannot. The predicted outcome of BayesR is hard to explain. It is not the risk of getting disease, since the estimated outcome can be bigger than 1.

When applying BayesRB and BayesR, convergence diagnostics should be conducted. Before applying the BayesRB or BayesR method, it is necessary to try different total numbers of MCMC loops, as well as numbers of burn-in loops, and select the ones where all the parameters mix well. But Moser et al. [2015] failed to do so. They used the default setting of the total number of MCMC loops and the number of burn-in loops of BayesR and they did not make diagnostic plots of their parameters, so it is not clear whether the parameters mix well in their MCMC process. In our studies, we conducted the convergence diagnostics in pilot simulated data analysis, genome-wide simulated data analysis and the real data analysis. Although, due to the limited computational time, BayesRB was run for less loops than needed to make all the parameters mix well, the convergence diagnostics still provide an idea of how well the parameters mixed.

To summarize, for SNP effect estimation, BayesRB has similar estimates to logistic regression for big effect SNPs, and shows BayesR’s sparseness characteristic for small effect SNPs. It makes up the defects of the conventional marginal logistic regression method and estimates the SNP effects taking account of the other SNPs. It also has better risk prediction performance than logistic regression and LASSO. Although BayesRB’s risk prediction performance is not better than BayesR and it does not have better associated SNP selection performance, BayesRB is still a promising method to use for dichotomous outcome data.

## 5 Software Availability

The BayesRB R package is available in the GitHub repository: https://github.com/sylviashanboo/BayesRB.

## 6 Acknowledgment

This study makes use of data generated by the Wellcome Trust Case-Control Consortium. A full list of the investigators who contributed to the generation of the data is available from www.wtccc.org.uk. Funding for the project was provided by the Wellcome Trust under award 076113, 085475 and 090355. The Wellcome Trust Case-Control Consortium and/or its Individual Investigators bear no responsibility for the further analysis or interpretation of these data, over and above that published by the Consortium.

This research was part of Dr. Shan’s dissertation (http://d-scholarship.pitt.edu/28629/) in the Department of Biostatistics, School of Public Health, University of Pittsburgh, Pittsburgh, Pennsylvania, United States of America.

## A Appendix: Detailed MCMC Steps in the BayesRB Algorithm

### Full conditional distribution for the general mean *μ* (MCMC step 2)

The logistic regression model can be written as a linear form with the latent variable *Z_i_*.

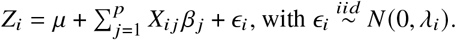

So, 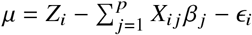

Then, 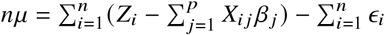, where 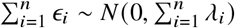.

Therefore, the conditional posterior distribution of the general mean *μ* is

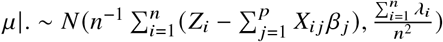

### Proof of the full conditional posterior distribution of *b_j_* and *β_j_* (MCMC step 3)

For SNP *j*, we update *β_j_* right after *b_j_*, and then we update them for SNP *j* + 1, which means we update *β_j_* and *b_j_* jointly.

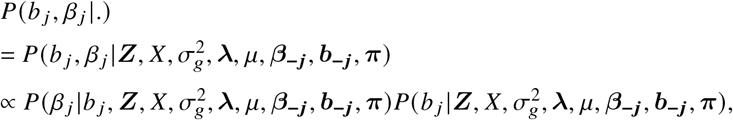

where ***b_−j_*** denotes the vector of categories that all the SNPs expect SNP *j* belong to. ***β_−j_*** denotes the vector of effects of all the SNPs expect SNP *j*.

1. First *b_j_* is updated by calculating 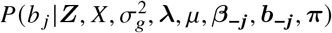:

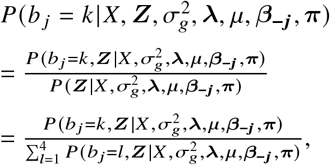

where *k* is the category SNP *j* is assigned to. *k* = (1, 2, 3, 4). Set 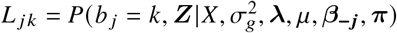, Then, 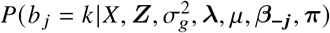

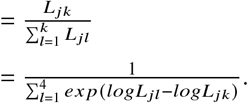 *L_jk_* is calculated below.

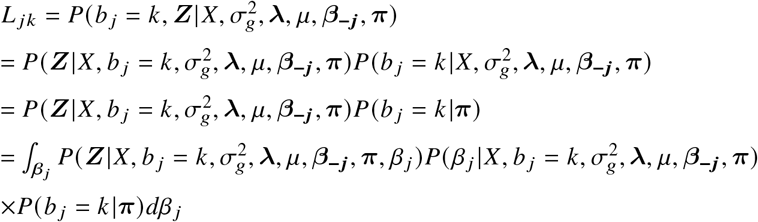 For individual *i*,

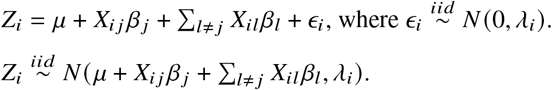 Therefore, 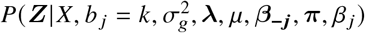

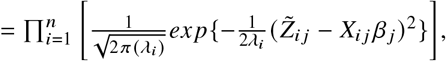

where 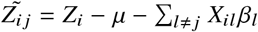. ***C*** is the vector (0, 10^−4^, 10^−3^, 10^−2^). Then,

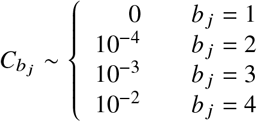 If *b_j_* ≠ 1,

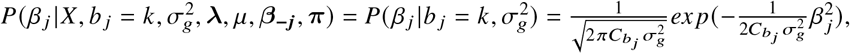

which is the prior of *β_j_* given *β_j_* is assigned to the category *b_j_*.

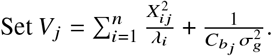 Then, 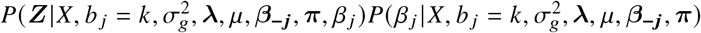

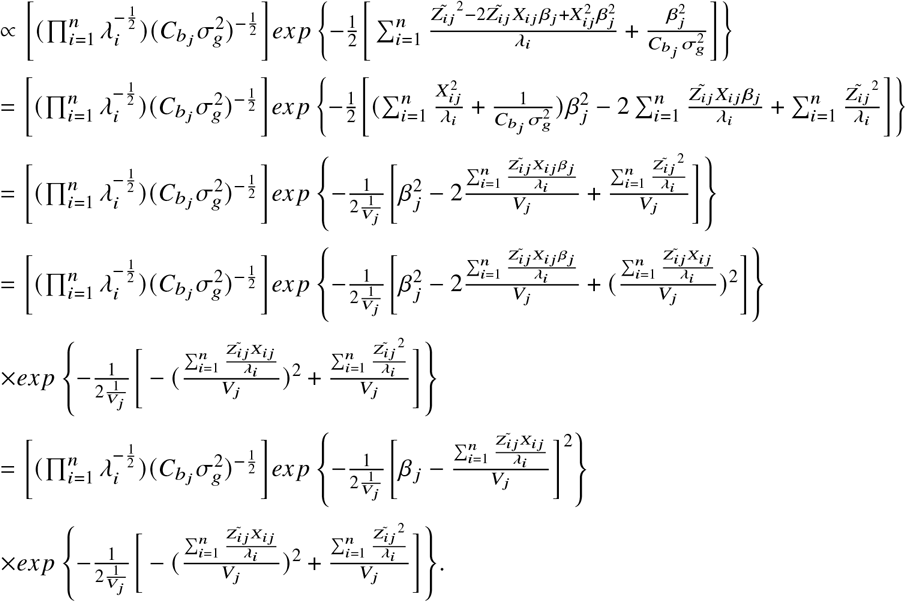 So, 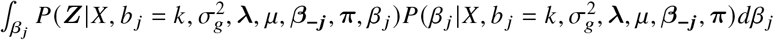,

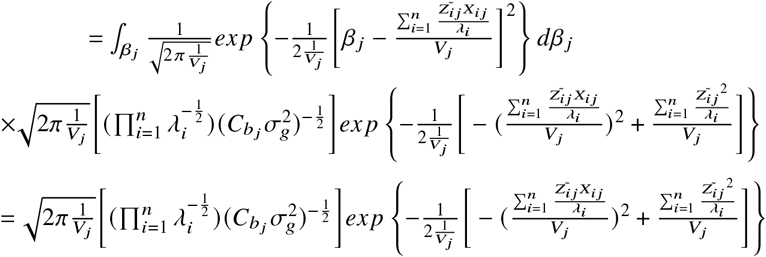 If *b_j_* ≠ 1,

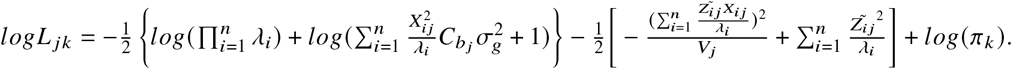 If *b_j_* = 1,

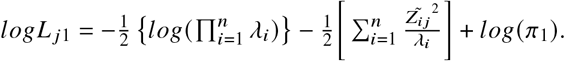 Set 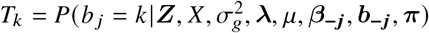 The SNP *j* is assigned to category *k* based on a value *h* sampled from a uniform distribution.

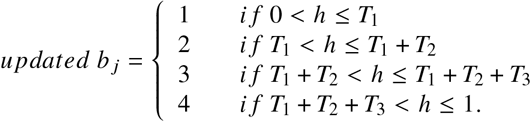
2. Then, we update *β_j_* by calculating 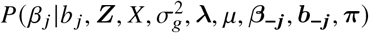

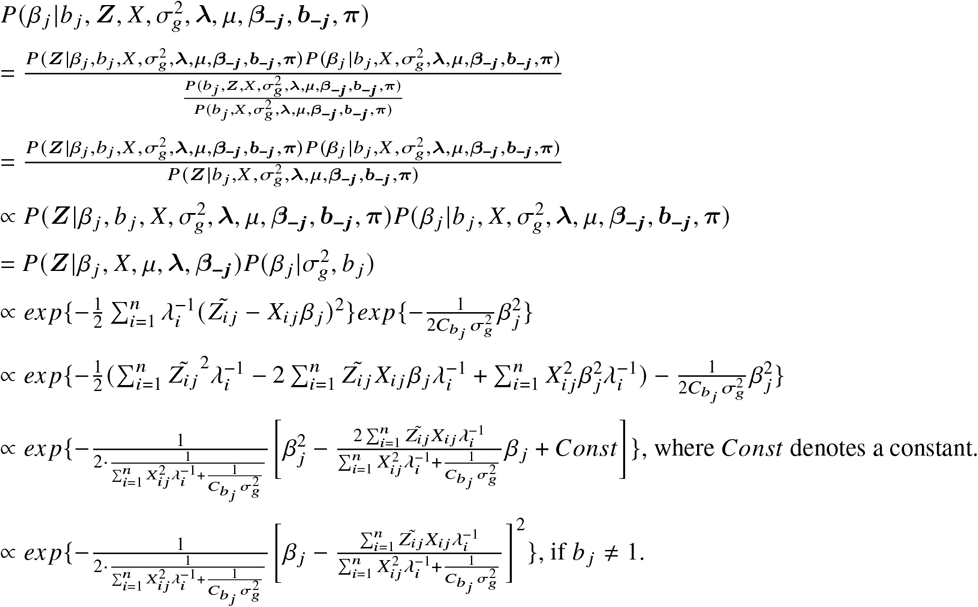 Therefore, the conditional posterior distribution of the effect of SNP *j*, which belongs to category *b_j_*, is

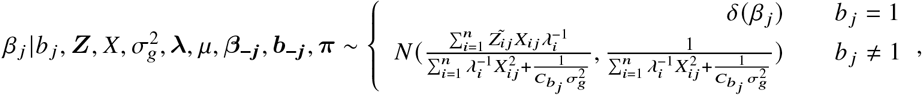

where *δ*(*β_j_*) denotes the dirac delta function with all probability mass at *β_j_* = 0 if *b_j_* = 1.

### Full conditional distribution for the relative variance for each mixture component 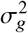 (MCMC step 4)

We assumed the SNPs are independent. From the dependency diagram,

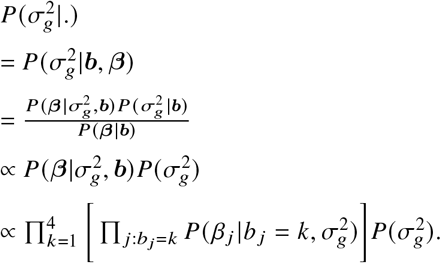

We use the uniform non-informative prior for 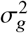. Then, 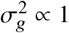.

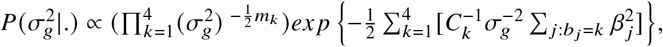

which is not a density function of any known distribution. Therefore, we used a Metropolis-Hasting sampling to update 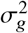. The initial value of 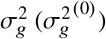 is sampled from N(0, U).

The steps are shown below:

1. Proposal function: 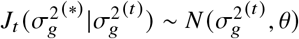 truncated at 0.

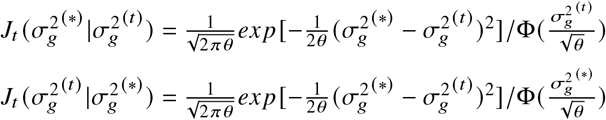
2. 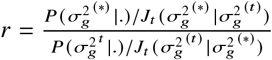

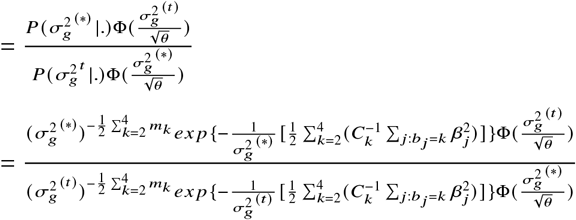
3. Sample *v* ~ *unif*(0, 1)
4. Update 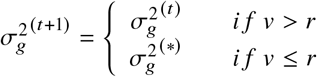 In the above steps, *U* and *θ* are tuned manually. Using the unrealistic data set, *U* = 200 and *θ* = 1.

## B Appendix: Pseudo Code of BayesRB

%% Set initial values

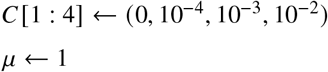

%% Initial value is when loop r=1

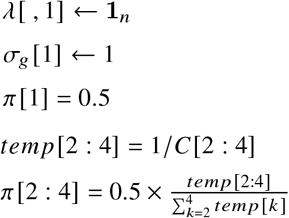

%% Initial value of *β_j_*’s are estimated from a regular marginal logistic regression.

%% Initial value of *μ* is estimated by taking the average of the intercepts in the marginal logistic regressions.

%% We record the initial value of *β_j_* and *μ* as *β* [*j*, *r*] and *μ* [*r*], where *r* = 1.

%% *β* [*j*, *r*] records the effect of SNP *j* in the *r*th iteration.

%% Initial value of 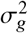 is sampled from a uniform distribution

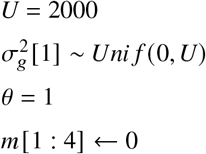

%% Finish setting the initial values

%% Start the MCMC steps:

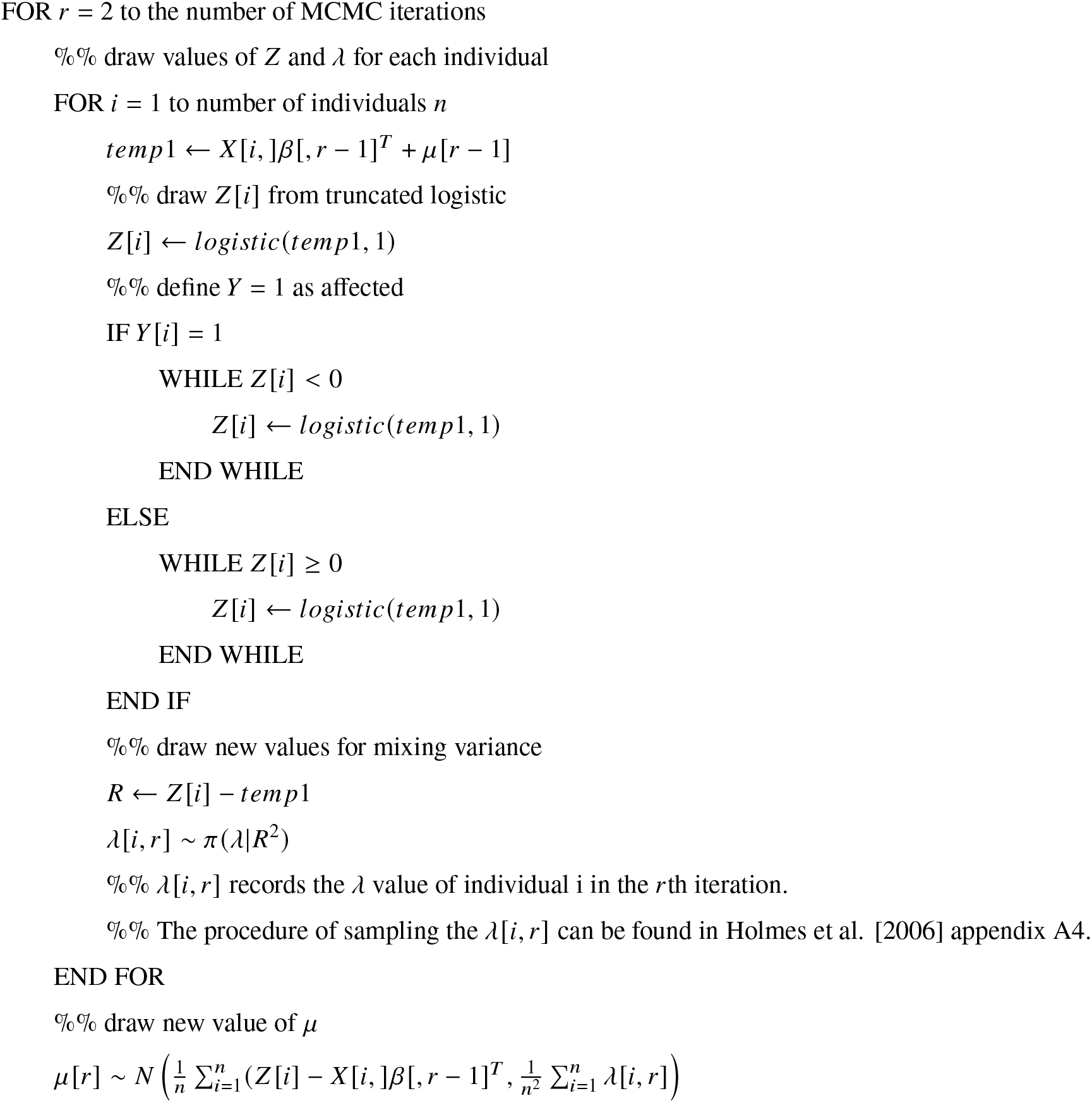
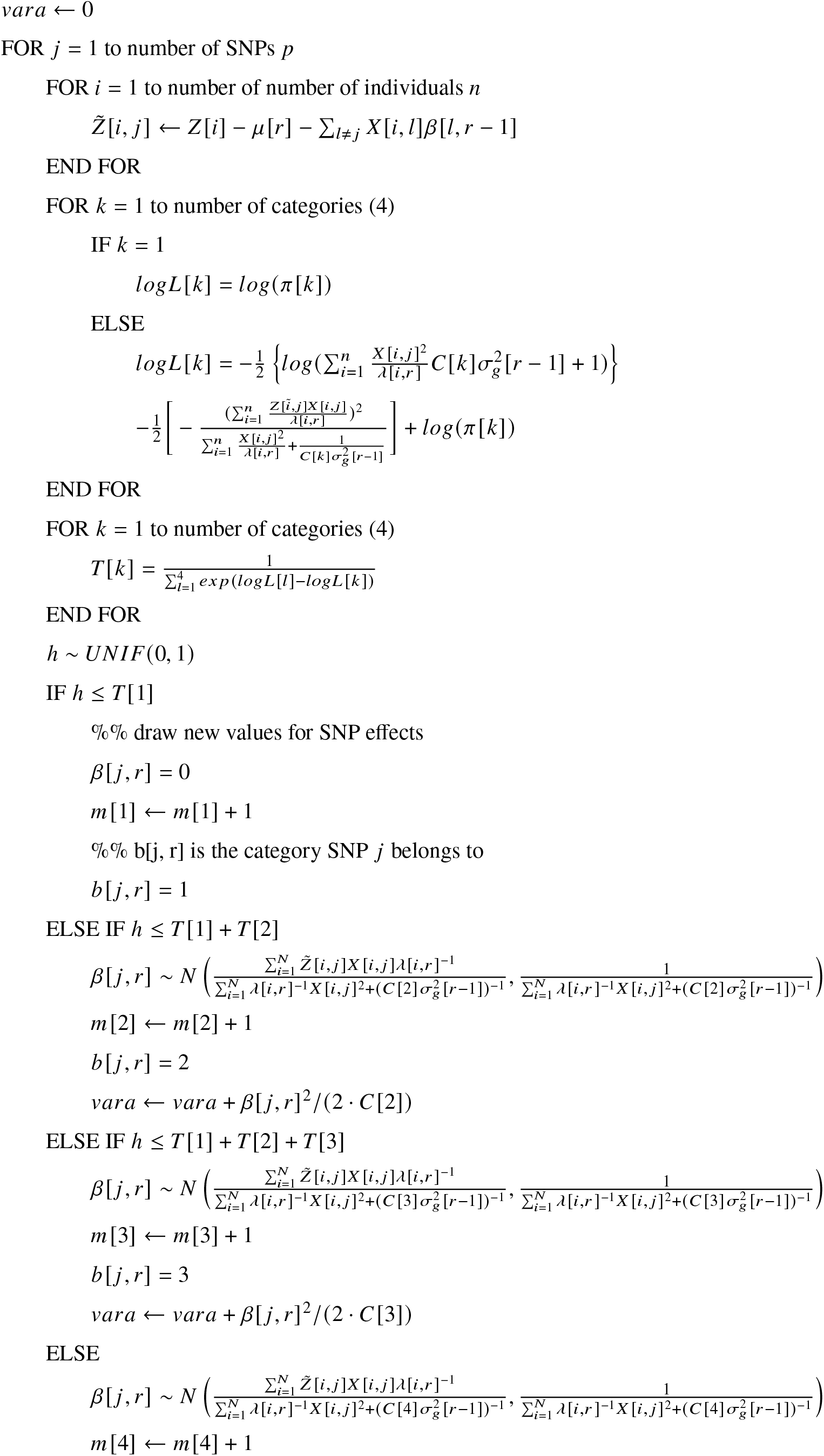
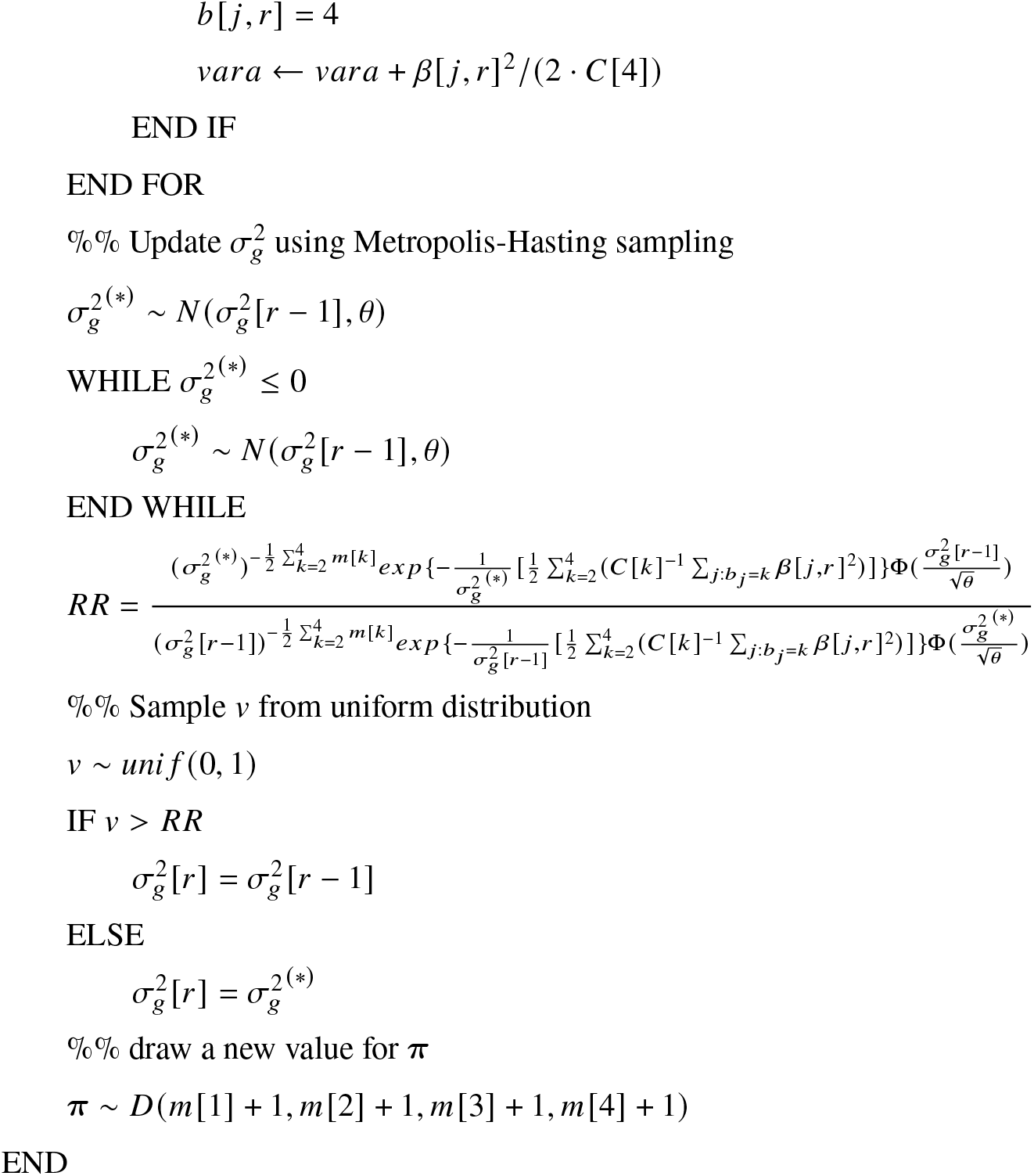

## Notes

### Competing Interest Statement

The authors have declared no competing interest.

